# FBXO47 is essential for preventing the synaptonemal complex from premature disassembly in mouse male meiosis

**DOI:** 10.1101/2021.11.17.469052

**Authors:** Nobuhiro Tanno, Kazumasa Takemoto, Yuki Takada-Horisawa, Ryuki Shimada, Sayoko Fujimura, Naoki Tani, Naoki Takeda, Kimi Araki, Kei-ichiro Ishiguro

**Author notes:** These authors equally contributed (Nobuhiro Tanno, Kazumasa Takemoto).

## Abstract

Meiotic prophase is a prolonged G2 phase that ensures the completion of numerous meiosis-specific chromosome events. During meiotic prophase, homologous chromosomes undergo synapsis to facilitate meiotic recombination yielding crossovers. It remains largely elusive how homolog synapsis is temporally maintained and destabilized during meiotic prophase. Here we show that FBXO47 is the stabilizer of synaptonemal complex during male meiotic prophase. Disruption of FBXO47 shows severe impact on homologous chromosome synapsis and DSB repair processes, leading to male infertility. Notably, in the absence of FBXO47, although once homologous chromosomes are synapsed, the synaptonemal complex is precociously disassembled before progressing beyond pachytene. Remarkably, *Fbxo47* KO spermatocytes remain in earlier stage of meiotic prophase and lack crossovers, despite apparently exhibiting diplotene-like chromosome morphology. We propose that FBXO47 functions independently of SCF E3 ligase, and plays a crucial role in preventing synaptonemal complex from premature disassembly during cell cycle progression of meiotic prophase.

## Introduction

Meiosis consists of a single DNA replication followed by two rounds of chromosome segregation, which halves the chromosome number to ultimately produce haploid gametes. During meiotic prophase I, sister chromatids are organized into proteinaceous structures, termed axial element (AE) or chromosome axis (Zickler and Kleckner, 2015). Homologous chromosomes (homologs) then undergo synapsis, which is promoted by the assembly of synaptonemal complex (SC) (Cahoon and Hawley, 2016). Homolog synapsis facilitates meiotic recombination yielding crossovers, a process that produces physical linkages called chiasmata between the homologs (Baudat et al., 2013) (Keeney et al., 2014). While homolog synapsis persists until meiotic recombination is completed during pachytene, it is dissolved upon diplotene-Metaphase I transition. Thus, homolog synapsis and de-synapsis is temporally regulated. However, it remains elusive how homolog synapsis is temporally maintained and destabilized during meiotic prophase.

SCF (SKP1–Cullin–F-box) E3 ubiquitin ligase is a key regulator of cell cycle (Deshaies, 1999) (Cardozo and Pagano, 2004). Accumulating lines of evidence suggest that SCF is involved in homolog synapsis in a wide variety of organisms. In mouse, homologous chromosomes showed premature desynapsis in *Skp1* conditional KO spermatocytes (Guan et al., 2020), suggesting that SCF is required for the maintenance of SC during male meiotic prophase. In *Drosophila* female, SkpA, a SKP1 homolog, is required for the assembly and/or the maintenance of SC (Barbosa et al., 2021). In budding yeast *Saccharomyces cerevisiae*, depletion of *Cdc53* that encodes Cullin resulted in defects in SC formation (Zhu et al., 2021). Thus, SCF is involved in the process of homolog synapsis during meiotic prophase in diverse organisms.

Fbox-domain containing proteins act as a substrate recognition subunit in SCF E3 ubiquitin ligase (Kipreos and Pagano, 2000) (Jin et al., 2004) (Reitsma et al., 2017). It has been shown that Fbox-domain containing proteins are involved in homolog synapsis in a wide variety of organisms. In rice plant *Oryza sativa*, mutants of MEIOTIC F-box *MOF* (He et al., 2016) and another Fbox *ZYGO1* (Zhang et al., 2017) showed defects in DNA double-strand break (DSB) repair and bouquet formation during meiotic prophase. In budding yeast, temperature sensitive mutant of *Cdc4* that encodes F-box protein showed defective SC formation and DSB repair (Zhu *et al*., 2021). In *Drosophila* female, depletion of Slmb (βTrcp) and CG6758 (Fbxo42) caused impaired assembly and/or premature disassembly of SC (Barbosa *et al*., 2021). Although the substrates are yet to be indentified in most of the cases, Fbox-domain containing proteins directly or indirectly regulate the assembly and disassembly of SC.

Previously, we identified MEIOSIN that plays an essential role in meiotic initiation both in mouse male and female (Ishiguro et al., 2020). MEIOSIN together with STRA8 (Kojima et al., 2019) activates meiotic genes and directs the switching from mitosis to meiosis. In the present study, we identified *Fbxo47* gene that encodes a Fbox protein, as one of the MEIOSIN/STRA8-target genes. Previous genetic studies suggested FBXO47 homologs are implicated in the progression of meiotic prophase in different species. In *C. elegance,* mutation in *prom-1* that encodes putative *Fbxo47* homolog, showed reduced homologous chromosome pairing and bivalent formation (Jantsch et al., 2007). In medaka fish, *fbxo47* mutant fails to complete meiotic prophase in female but switches developmental fate from oogenesis into spermatogenesis (Kikuchi et al., 2020). In mouse, *Fbxo47* gene that has previously been identified as a meiotic gene by single cell RNA-seq analysis of testes, is essential for mouse spermatogenesis (Chen et al., 2018) (Hua et al., 2019). Although previous studies suggest that FBXO47 homologs and distant meiotic Fbox-domain containing proteins play a role in homologous chromosome pairing/synapsis and meiotic recombination in a wide variety of organisms, the precise mechanisms how these proteins are involved in these processes remained elusive. Furthermore, whether FBXO47 is indeed involved in the function of SCF is unknown. Here we show that mouse FBXO47 is essential for mainatining homolog synapsis during meiotic prophase. FBXO47 is a cytoplasmic protein rather than a telomere binding protein, and functions independently of SCF. We demonstrate that in *Fbxo47* KO spermatocytes, homologous chromosome synapsis is complete, but SC is precociously disassembled. Further, we show that *Fbxo47* KO spermatocytes fail to progress beyond pachytene and remain in earlier meiotic prophase in terms of cell cycle progression, despite the apparent exhibition of diplotene-like morphology of chromosomes. We propose that FBXO47 is essential for preventing SC from premature destruction during cell cycle progression of male meiotic prophase. Further, we discuss the different observations and interpretations between the present study and the previous study on FBXO47 (Hua *et al*., 2019).

## Results

### FBXO47 is expressed in mouse testes

Previously, we demonstrated that MEIOSIN collaborating with STRA8 activates meiotic genes, which are required for numerous meiotic events (Ishiguro *et al*., 2020). In spermatocytes, we identified *Fbxo47* as one of the MEIOSIN/STRA8-bound genes (Fig. 1A). Our previous RNA-seq analysis showed that expression of *Fbxo47* was significantly downregulated in *Meiosin* KO testes at postnatal day 10 (P10) when a cohort of spermatocytes should undergo the first wave of meiotic entry (Ishiguro *et al*., 2020). We confirmed this by RT-qPCR analysis demonstrating that *Fbxo47* expression level was indeed downregulated in *Meiosin* KO testis at P10 (Fig. 1B). We further examined the expression patterns of *Fbxo47* in different mouse tissues by RT-PCR analysis. *Fbxo47* gene showed higher expression levels in adult testis compared to other adult organs that we examined (Fig. 1C). Spermatogenic expression of *Fbxo47* gene was further confirmed by the reanalysis of previous scRNA-seq data of adult mouse testis (Hermann et al., 2018) (Fig. 1D). The result indicated that *Fbxo47* was coordinately expressed with the landmark genes of meiotic spermatocyte such as *Dmc1*, and spermatid at spermiogenesis such as *Acrv1*, rather than those of spermatogonia such as *Zbtb16* (Fig. 1D). We noticed that *Fbxo47* mRNA was expressed weakly in meiotic spermatocytes, and highly in spermatids in testes, which is consistent with a previous study (Chen *et al*., 2018). In females, expression of *Fbxo47* mRNA was examined by the reanalysis of previous scRNA-seq data of fetal ovaries (Shimada et al., 2021). We found that *Fbxo47* was coordinately expressed during meiotic prophase, such as *Dmc1* (Fig. 1E). Expression of *Fbxo47* mRNA culminated at E16.5 and declined afterward in the ovary (Fig. 1F).

**Figure 1.**
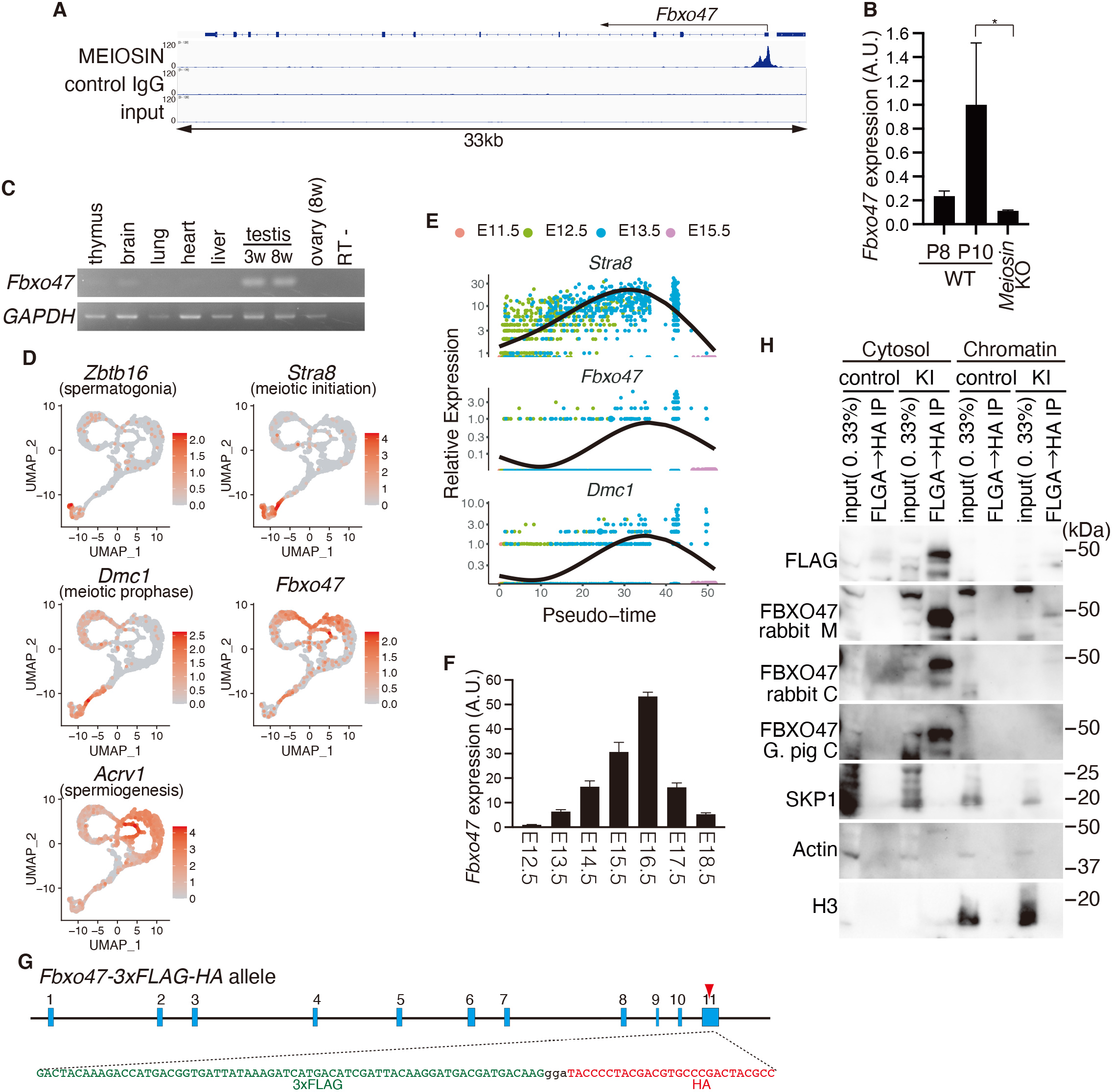
Identification of the meiosis-specific factor FBXO47. **(A)** Genomic view of MEIOSIN binding peak over *Fbxo47* loci. Genomic coordinates were obtained from Ensembl. **(B)** The expression of *Fbxo47* in WT and *Meiosin* KO was examined using RT-PCR. Testis RNA was obtained from WT (3 animals each for P8 and P10) and *Meiosin* KO (3 animals). The graph shows the expression level of *Fbxo47* normalized by that of *GAPDH* with SD. Expression level of *Fbxo47* in P10 WT was set to 1. Statistical significance is shown by *p*-value (Two-tailed t-test). *: p < 0.05. **(C)** The tissue-specific expression pattern of *Fbxo47* was examined by RT-PCR. Testis RNA was obtained from embryonic day 18 (E18), 3-weeks old (3w) and 8-weeks old (8w) male mice. Ovary RNA was obtained from adult 8-weeks old (8w) female mice. RT-indicates control PCR without reverse transcription. **(D)** Expression patterns of *Fbxo47* and other key developmental genes are reanalyzed using public scRNA-seq data of spermatogenic cells in adult mouse testis (GEO: GSE109033) (Hermann *et al*., 2018). Expression patterns of *Fbxo47* and other key developmental genes are shown in UMAP plots. Key developmental genes include *Zbtb16*: spermatogonia, *Stra8*: differentiating spermatogonia and pleleptotene spermatocyte, *Dmc1*: meiotic prophase spermatocyte, *Acrv1*: round and elongated spermatid. UMAP of *Zbtb16 and Stra8* was adopted from our previous study (Horisawa-Takada et al., 2021). **(E)** Expression profiles of *Fbxo47, Stra8* and *Dmc1* in E11.5, E12.5, E13.5, E15.5 fetal ovaries along pseudotime trajectory of germ cells. Pseudotime analysis was performed by reanalyzing scRNA-seq data (DRA011172) (Shimada *et al*., 2021). Pseudotime expression profile of *Stra8* was adopted from our previous study (Horisawa-Takada *et al*., 2021). **(F)** The expression pattern of *Fbxo47* in the embryonic ovary was examined by RT-qPCR. Average values normalized to E12.5 gonad are shown with SD from technical triplicates or quadruplicates. N=1 gonadal sample for each embryo. **(G)** Schematic illustrations of the *Fbxo47-3xFLAG-HA* knock-in (*Fbxo47-3FH* KI) allele. Blue boxes represent exons. The stop codon in the exon 11was replaced with in-frame *3xFLAG-HA* and the endogenous 3’UTR. **(H)** Western blot showed immunoprecipitates after tandem affinity purifications using anti-FLAG and anti-HA from cytoplasmic and chromatin extracts of WT (non-tagged control) and *Fbxo47-3FH* KI mouse testes (P15-18). The same membrane was sequentially reblotted with different antibodies against the endogenous FBXO47 that we generated. rabbit M: rabbit anti-FBXO47 middle region, rabbit C: rabbit anti-FBXO47 C-terminal region, G.pig C: guinia pig anti-FBXO47 C-terminal region.

To determine the meiotic stage-specific expression of FBXO47 protein, we generated different antibodies against FBXO47 C-terminal region (aa 271-451) and middle region (aa173-316). However, we failed to evaluate stage specificity of endogenous FBXO47 protein expression by immunostaining, although it was uncertain whether this was due to the sensitivity of the antibodies, inaccessibility of the antibodies to the epitopes, or low expression level of FBXO47 protein in the target cells.

To circumvent this issue, we generated *Fbxo47*-*3xFLAG-HA* knock-in (*Fbxo47*-*3FH-GFP* KI) mice, which allowed the detection of FBXO47-3xFLAG-HA protein esxpressed from endogenous *Fbxo47* locus (Fig. 1G, Fig. S1). We examined FBXO47-3xFLAG-HA fusion protein from cytosolic and chromatin extracts of *Fbxo47*-*3FH-GFP* KI testes. Immunoblotting demonstrated that FBXO47 protein was detected with FLAG antibody only when it was enriched by tandem immunoprecipitations using anti-FLAG and anti-HA antibodies (Fig. 1H), suggesting that the expression level of FBXO47 protein was low in testes. We noticed that more FBXO47 protein was detected in the cytosolic fraction compared to the chromatin fraction (Fig. 1H), suggesting its predominant locatlization in the cytoplasm rather than on the chromatin. Sequential reblotting showed that different antibodies against the endogenous FBXO47 protein that we generated detected the same protein as indicated by anti-FLAG antibody (Fig. 1H). Previous study showed that FBXO47 binds to telomeric proteins TRF1 and TRF2 (Hua *et al*., 2019). However, we failed to detect neither TRF1 nor TRF2 in FBXO47 immunoprecipitates from testis chromatin fraction (Fig. S3A), which was a sharp contrast to the previous study (Hua *et al*., 2019).

### FBXO47 may function independently of SCF in spermatocytes

FBXO47 possesses a putative Fbox domain, whose biological function has remained elusive. It is well known that Fbox-domain containing proteins confers substrate specificity to SCF (SKP1–Cullin–F-box) E3 ubiquitin ligase (Jin *et al*., 2004), and 69 different Fbox proteins are estimated to be encoded in human genome (Reitsma *et al*., 2017). This prompted us to examine whether SKP1, a major core subunit of SCF, was co-immunoprecipitaed with FBXO47 by immunoblot and mass spectrometry analysis (Fig. 1H, Fig. S2). However, we failed to detect SKP1 in FBXO47 immunoprecipitates.

To further examine whether FBXO47 serves as a subunit of SCF by reciprocal immunoprecipitation of SKP1, we generated *Skp1*-*3xFLAG-HA* knock-in (*Skp1*-*3FH-GFP* KI) mice, which allowed the detection of SKP1-3xFLAG-HA protein esxpressed from endogenous *Skp1* locus and its associated factors (Fig. 2A). Although the homozygous *Skp1-3xFLAG-HA* KI mice were embryonic lethal, heterozygous knock-in mice were fertile and developed normally. Consistent with a previous study (Guan *et al*., 2020), SKP1-3xFALG-HA fusion protein localized along the SC in the *Skp1*-*3FH-GFP* KI spermatocytes (Fig. 2B). SKP1-3xFLAG-HA was enriched by tandem immunoprecipitations using anti-FLAG and anti-HA antibodies from testis cytosolic fraction (Fig. 2C). Mass spectrometry analysis demonstarated that total of 45 different Fbox-domain containing proteins and SCF core subunits (SKP1, RBX1, CUL1, CUL7) were co-immunoprecipitated with SKP1-3xFLAG-HA (Fig. 2D, Supplementary Data1).

**Figure 2.**
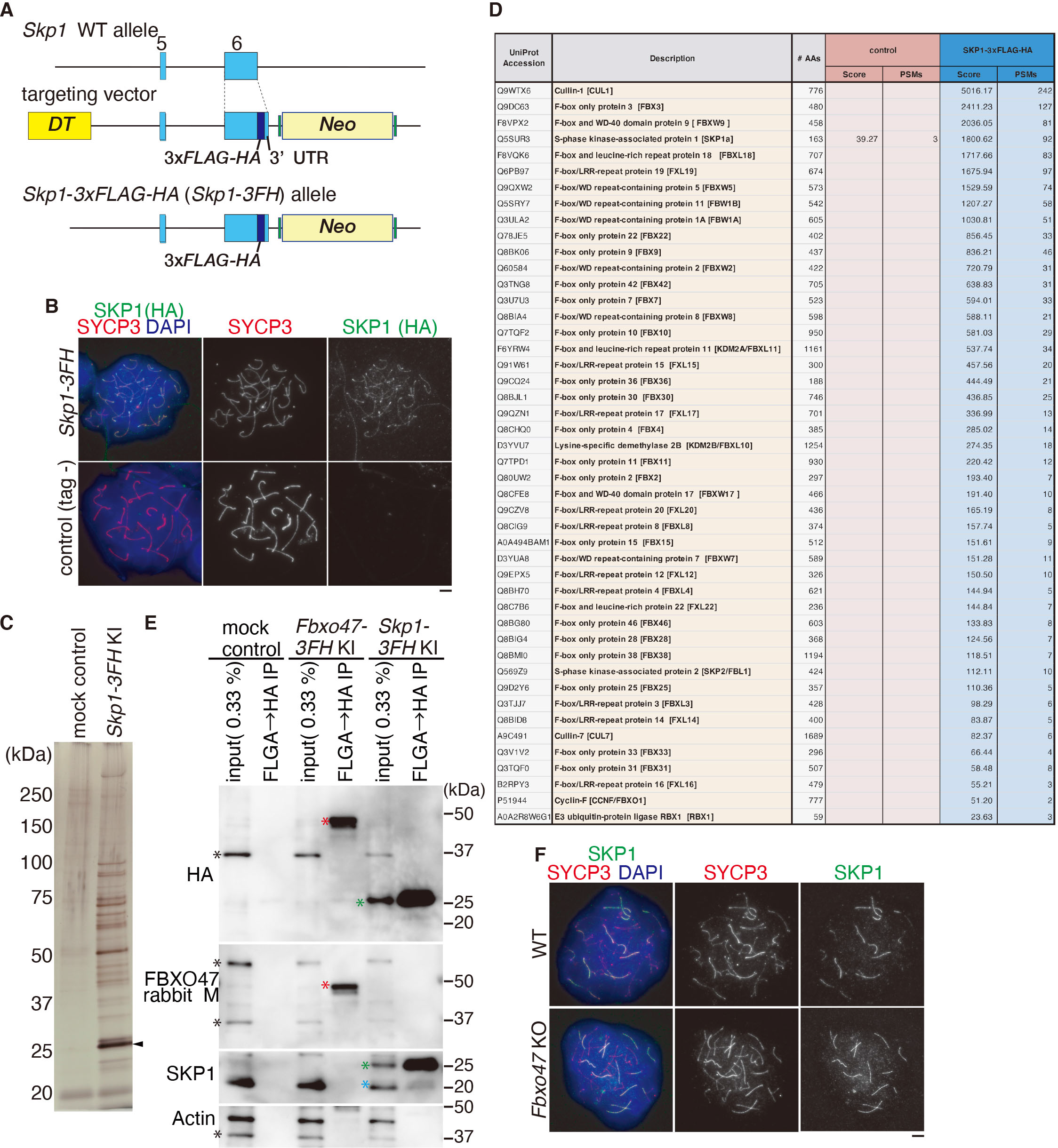
FBXO47 is not involved in the function of SCF. **(A)** Schematic illustrations of the *Skp1-3xFLAG-HA* knock-in (*Skp1-3FH* KI) allele. Blue boxes represent exons. The stop codon in the exon 6 was replaced with in-frame *3xFLAG-HA* and the endogenous 3’UTR. **(B)** Chromosome spreads of WT (non-tagged) and *Skp1-3FH* KI spermatocytes were immunostained as indicated. Scale bar: 5 μ **(C)** Silver staining of the immunoprecipitates from cytosolic extracts of WT (non-tagged control) and *Skp1-3FH* KI mouse testes after tandem affinity purifications using anti-FLAG and anti-HA antibodies. Arrowhead: SKP1-3xFLAG-HA. **(D)** The immunoprecipitates from the cytosolic fraction of the WT (non-tagged control) and *Skp1-3FH* KI testis extracts were subjected to liquid chromatography tandem-mass spectrometry (LC-MS/MS) analyses. The Fbox-containing proteins and SCF subunits identified by the LC-MS/MS analysis are presented after excluding the proteins detected in the control mock purification. The proteins are listed with SwissProt accession number, the number of peptide hits and Mascot scores. Full list of identified proteins are shown in the Supplementary Data1. It is worth noting that SC central element components, Six6OS1 and SYCE1, were included in the LC-MS/MS data of SKP1-3xFLAG-HA immunoprecipitates (Supplementary Data1). This suggests that SCF E3 ubiquitin ligese may target those SC components using an F-box protein listed in the LC-MS/MS data as a substrate recognition subunit. **(E)** Western blot showed immunoprecipitates from cytosolic extracts of WT (non-tagged control), *Fbxo47-3FH* KI and *Skp1-3FH* KI (heterozygous) testes after tandem affinity purifications using anti-FLAG and anti-HA antibodies. The same membrane was sequentially reblotted with different antibodies as indicated. Red *: FBXO47-3xFLAG-HA, Green *: SKP1-3xFLAG-HA, Blue *: endogenous SKP1, Black *: non-specific band. Note that SKP1 was not detected in FBXO47 immunoprecipitate from *Fbxo47-3FH* KI testis extracts, and reciprocally FBXO47 was not detected in SKP1 immunoprecipitate from *Skp1-3FH* KI testis extracts. **(F)** Chromosome spreads of WT and *Fbxo47* KO spermatocytes were immunostained as indicated. Scale bar: 5 μm.

However, we failed to detect FBXO47 in the SKP1-3xFLAG-HA immunoprecipitates either by mass spectrometry analysis or by western blotting (Fig. 2D, E). SKP1 localized along the SC in *Fbxo47* KO, suggesting that localization of SKP1 did not depend on FBXO47 (Fig. 2F). Altogether, our data suggest that FBXO47 may function independently of SCF in mouse testes. Previous study showed that FBXO47 interacts with SKP1 in yeast two hybrid assay and in GFP-SKP1 IP using HEK293T cell extract that overexpressed FLAG-FBXO47 and GFP-SKP1(Hua *et al*., 2019). Although we do not know the exact reason for these controversial observations between our present study and the previous one (Hua *et al*., 2019), this could be due to their detection methodology using yeast and overexpression of FBXO47 in culuture cells.

### Expression of FBXO47 is limitted to early meiotic prophase in mouse testes

To identify the specific stage in which FBXO47 was expressed, we performed immunostaining using stage specifc markers SYCP3 (a component of meiotic chromosome axis), SYCP1 (a marker of homologous chromosome synapsis), and γH2AX (a marker of DSBs). Immunostaining of the *Fbxo47*-*3FH-GFP* KI testis (P15) indicated that FBXO47 protein was detected by HA antibody in average 21% (n = 3) among total SYCP3 positive seminiferous tubules (Fig. 3A). Close inspection of seminiferous tubules showed that FBXO47 protein indicated by the presence of HA staining appeared in the cytosol at leptotene and zygotene (Fig. 3B, C). Notably, the expression level of FBXO47-3xFLAG-HA fusion protein declined in pachyetene, when homologs were fully synapsed (Fig. 3C). Testis-specific histone H1t is a marker of spermatocytes later than mid pachytene (Cobb et al., 1999) (Drabent et al., 1996). Immunostaining of seminiferous tubules by testis-specific histone H1t indicated that FBXO47 protein was expressed only in H1t negative stage (Fig. 3D). None of H1t positive spermatocytes showed FBXO47 immunostaining (Fig. 3E), suggestiung that FBXO47 expression had declined by mid-pachytene. Thus, the expression of FBXO47 protein was limited to a narrow window of early meiotic prophase. Although the expression of *Fbxo47* mRNA was upregulated in spermatids, immunostaining of FBXO47 protein detected no more than background levels in spermatids (Fig. 3F). This suggested that the expression of FBXO47 might be post-transcriptionally suppressed after post-meiotic spermatids to have the expression specifically limited to early meiotic prophase.

**Figure 3.**
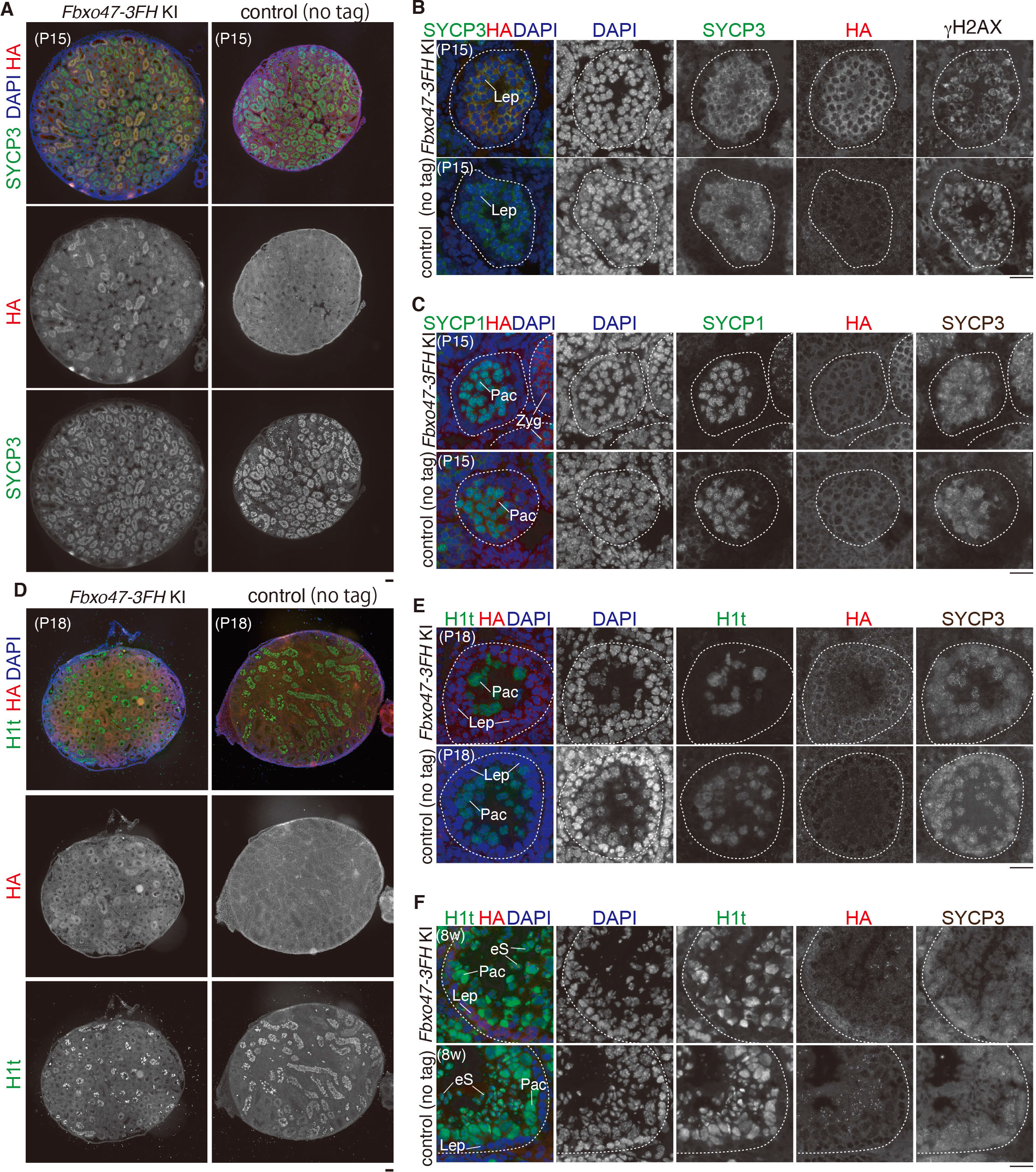
FBXO47 was expressed in early meiotic prophase. **(A)** Testis sections from *Fbxo47-3FH* KI and control (non-tagged) mice (P15) were stained for HA, SYCP3 and DAPI. Average 21% of the seminiferous tubules that have SYCP3+spermatocytes showed HA+/SYCP3+ in *Fbxo47-3FH* KI testes (n = 3 animals), while none of those was HA+/SYCP3+ in WT (n = 3 animals). Scale bar: 100 μm. **(B)** Seminiferous tubule sections from *Fbxo47-3FH* KI and control (non-tagged) mice (P15) were stained for HA, SYCP3, γH2AX and DAPI. Lep: leptotene. Scale bar: 25 μm. **(C)** Seminiferous tubule sections were stained for HA, SYCP3, SYCP1 and DAPI as in (B). Zyg: zygotene, Pac: pachytene spermatocyte, rS: round spermatid, eS: elongating spermatid. Scale bar: 25 μm. **(D)** Testis sections from *Fbxo47-3FH* KI and control (non-tagged) mice (n = 3 for each genotype, P18) were stained for HA, H1t and DAPI as in (A). Number of seminiferous tubules that have HA+/ H1t+ cells was counted per the seminiferous tubules that have H1t+ spermatocyte cells (52, 36, 18 tubules for non-tagged control; 15, 51, 36 tubules for *Fbxo47-3FH* KI mice). Scale bar: 100 μm. **(E)** Seminiferous tubule sections (P18) were stained for HA, SYCP3, H1t and DAPI as in (B). Lep: leptotene, Pac: pachytene spermatocyte, rS: round spermatid, eS: elongating spermatid. Scale bar: 25 μm. **(F)** Seminiferous tubule sections (8-weeks old) were immunostained as in (E). Lep: leptotene, Pac: pachytene spermatocyte, eS: elongating spermatid. Scale bar: 25 μm. Note that pachy signals of HA immunostaining were nonspecific, since they were visible in control.

### Disruption of *Fbxo47* led to severe defect in spermatogenesis

In order to address the role of *Fbxo47* in meiosis, we deleted Exon3-Exon11 of *Fbxo47* loci in C57BL/6 fertilized eggs through the CRISPR/Cas9 system (Fig. 4A). RT-PCR analysis showed that *Fbxo47* mRNA expression level was absent in *Fbxo47* KO testis (Fig. 4B). Although *Fbxo47* KO male mice did not show overt phenotype in somatic tissues, defects in male reproductive organs were evident with smaller-than-normal testes (Fig. 4C). Histological analysis revealed that post-meiotic spermatids and spermatozoa were absent in eight-week-old *Fbxo47* KO seminiferous tubules (Fig. 4D). Accordingly, sperm was absent in adult *Fbxo47* KO caudal epididymis (Fig. 4E). Consistently, seminiferous tubules that contain PNA lectin (a marker of spermatids) positive cells were absent in *Fbxo47* KO (Fig. 4F). Thus, the later stage of spermatogenesis was severely abolished in *Fbxo47* KO seminiferous tubules, resulting in male infertility (Fig. 4G). In contrast to male, *Fbxo47* KO females exhibited seemingly normal fertility with no apparent defects in adult ovaries (Fig. 4H). Consistent with this histological observation of ovaries, metaphase I oocytes derived from *Fbxo47* KO females processes normal number of bivalent chromosomes with chiasmata, indicating that *Fbxo47* KO oocytes had progressed normal meiotic prophase (Fig. 4I). Furthermore, *Fbxo47* KO females were fertile (Fig. 4G, J), although we could not exclude the possibility that more subtle defects might have occurred in the ovaries besides fertility. Thus, the infertility caused by disruption of *Fbxo47* was male specific. Therefore, these results suggest that requirement of FBXO47 is sexually different in mouse.

**Figure 4.**
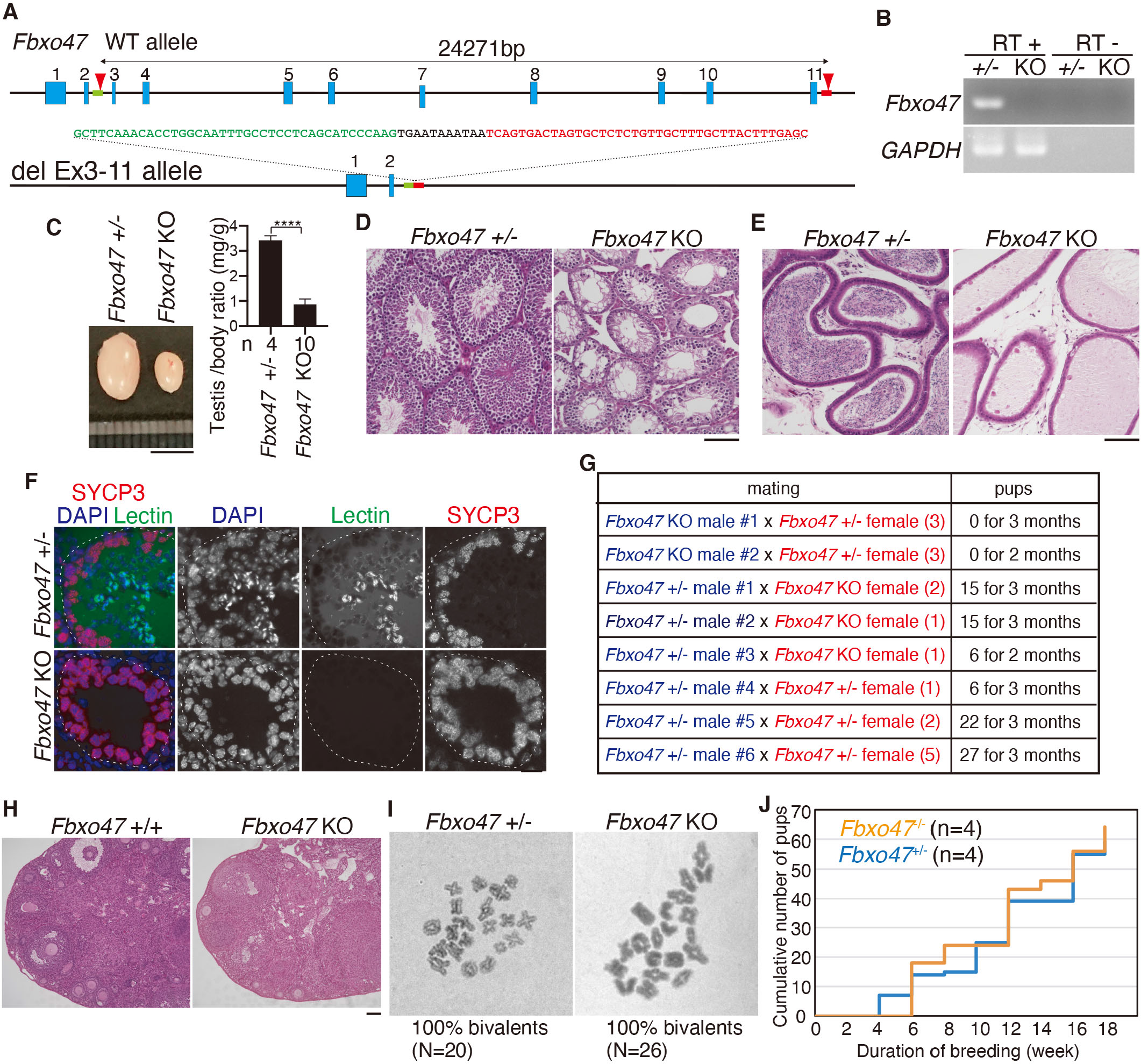
Spermatogenesis was impaired in *Fbxo47* knockout male. **(A)** The allele with targeted deletion of Exon3-13 in *Fbxo47* gene was generated by the introduction of CAS9, the synthetic gRNAs designed to target intron2 and the downstream of Exon11 (arrowheads), and ssODN (green and red boxes) into C57BL/6 fertilized eggs. **(B)** *Fbxo47* mRNA expression was examined by RT-PCR. Testis RNA was obtained from *Fbxo47*+/- and *Fbxo47* KO males (P13). RT-indicates control PCR without reverse transcription. **(C)** Testes from *Fbxo47*+/- and *Fbxo47* KO (8-weeks old). Testis/body-weight ratio (mg/g) of *Fbxo47*+/- and *Fbxo47* KO mice (8-weeks old) is shown on the right (Mean with SD). n: the number of animals examined. Statistical significance is shown by ****: *p* < 0.0001 (Two-tailed t-test). Scale bar: 5 mm. **(D)** Hematoxylin and eosin staining of the sections from *Fbxo47*+/- and *Fbxo47* KO testes (8-weeks old). Biologically independent mice for each genotype were examined. Scale bar: 100 μm. **(E)** Hematoxylin and eosin staining of the sections from *Fbxo47*+/- and *Fbxo47* KO epididymis (8-weeks old). Biologically independent mice for each genotype were examined. Scale bar: 100 μm. **(F)** Seminiferous tubule sections (8-weeks old) were stained for SYCP3, PNA lectin and DAPI. Note that the seminiferous tubule that contained PNA-positive elongated spermatids were not identified in *Fbxo47* KO testes. Scale bar: 25 μm. **(G)** Number of pups born by mating *Fbxo47*+/- and *Fbxo47* KO males with *Fbxo47*+/- or *Fbxo47* KO females (N = number of females in the same cage) to examine fertility. *Fbxo47* KO male #1 was initially mated with three *Fbxo47*+/- females (all 6-weeks old at the start point of mating). After one month, another *Fbxo47* KO male #2 was started to cohabit with those females (8-weeks old at the start point of mating). This cage was observed for 3 months from the start of mating. **(H)** Hematoxylin and Eosin stained sections of *Fbxo47*+/- and *Fbxo47* KO ovaries (8-weeks old). Scale bar: 100 μm. **(I)** Giemza staining of metaphase I chromosomes from *Fbxo47*+/- (N=20) and *Fbxo47* KO spermatocytes (N=26). **(J)** Cumulative number of pups born from *Fbxo47*+/- (n=4, all 6-weeks old at the start point of mating) and *Fbxo47* KO (n=4, all 6-weeks old at the start point of mating) females.

### Synaptonemal complex was prematurely disassembled in *Fbxo47* KO spermatocytes

To further investigate at which stage the primary defect appeared in the *Fbxo47* KO, we analyzed the progression of spermatogenesis by immunostaining. Testis-specific histone H1t is a marker of spermatocytes later than mid pachytene and round spermatids (Cobb *et al*., 1999) (Drabent *et al*., 1996). Close inspection of the seminiferous tubules (3 week) by immunostaining with antibodies against H1t along with SYCP3 (a component of meiotic chromosome axis) indicated that *Fbxo47* KO spermatocytes failed to reach mid pachytene, whereas spermatocytes in age-matched control passed beyond mid pachytene as indicated by the presence of H1t staining (Fig. 5A). This suggests that progression of meiotic prophase was blocked in *Fbxo47* KO spermatocytes. Immunostaining analysis of spread chromosome with antibodies against SYCP3 along with SYCP1 (a marker of homolog synapsis) demonstrated that *Fbxo47* KO spermatocytes underwent homologous chromosome synapsis and seemingly reached pachytene stage as in age-matched control (Fig. 5B).

**Figure 5.**
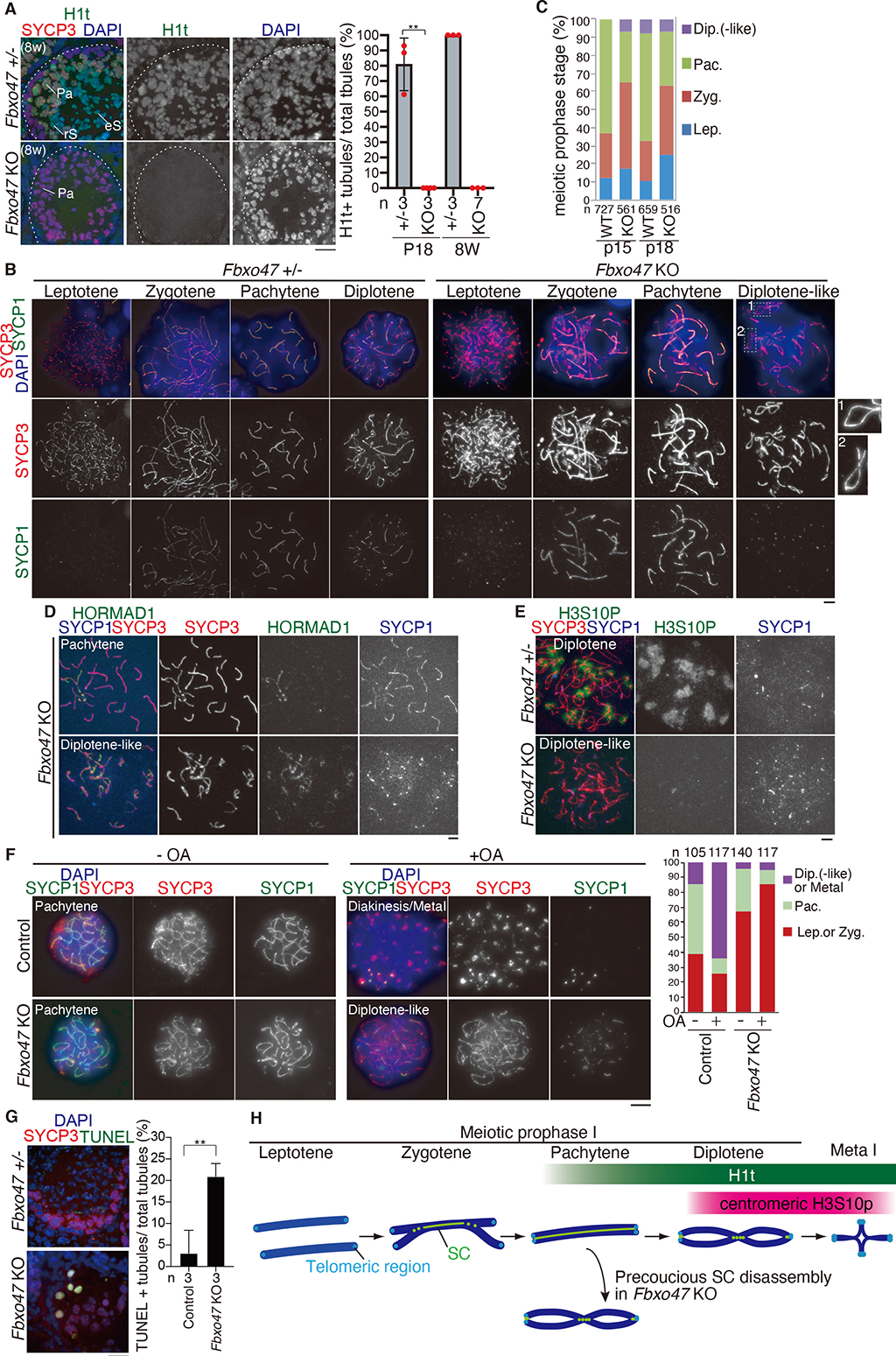
Premature disassembly of SC in *Fbxo47* KO spermatocytes. **(A)** Seminiferous tubule sections (P18 and 8-weeks old) were stained for SYCP3, H1t and DAPI. Pa: pachytene spermatocyte, rS: round spermatid, eS: elongating spermatid. Shown on the right is the quantification of the seminiferous tubules that have H1t+/SYCP3+ cells per the seminiferous tubules that have SYCP3+ spermatocyte cells in WT and *Fbxo47* KO mice (Mean with SD). n: the number of animals examined for each genotype. Statistical significance is shown (Unpaired t-test). ** : *p* = 0.0012 for *Fbxo47* heterozygous versus *Fbxo47* KO at P18. *Fbxo47* heterozygous (p18: 62, 61, 29 tubules/animal were counted from 3 animals; 8w: 135, 143, 45 tubules/animal were counted from 3 animals) and *Fbxo47* KO testes (p18: 105, 59, 141 tubules/animal were counted from 3 animals; 8w: 36, 55, 63, 64, 88, 69, 108 tubules/animal were counted from 7 animals). Scale bar: 25 μm. **(B)** Chromosome spreads of WT and *Fbxo47* KO spermatocytes (3-4 weeks old) were immunostained as indicated. Enlarged images are shown to highlight de-synapsed chromosomes in diplotene-like *Fbxo47* KO spermatocytes. Scale bar: 5 μm. **(C)** Quantification of meiotic prophase stage spermatocytes per total SYCP3+ spermatocytes in WT and *Fbxo47* KO mice at P15 and P18 is shown. n: the number of cells examined. **(D)** Chromosome spreads of pachytene and diplotene-like *Fbxo47* KO spermatocyte were immunostained for SYCP3, H3S10P and HORMAD1. Scale bar: 5 μm. **(E)** Chromosome spreads of diplotene spermatocyte in the control and diplotene-like spermatocyte in *Fbxo47* KO spermatocytes (P18) were immunostained for SYCP3, H3S10P and DAPI. Scale bar: 5 μm. Note that centromeric regions are positively stained for H3S10P in the control diplotene spermatocyte but not in diplotene-like spermatocyte in *Fbxo47* KO spermatocytes. **(F)** Spermatocytes isolated from the control *Fbxo47*+/- and *Fbxo47* KO testes were cultured *in vitro* in the presence or absence of OA for 3 hours. Quantification of meiotic prophase stage is shown on the right. n: the number of cells examined. Note that the control spermatocytes showed a typical feature of diakinesis/Meta I with condensed chromosomes and remaining SYCP3 at centromeres. **(G)** Seminiferous tubule sections from 8-weeks old mice were subjected to TUNEL assay with immunostaining for SYCP3. L: leptotene, Pa: pachytene. Shown on the right is the quantification of the seminiferous tubules that have TUNEL+ cells per total tubules in *Fbxo47*+/- (8w; n=3) and *Fbxo47* KO (8w; n=3) testes (mean with SD). Statistical significance is shown by ** *p* = 0.0072 (Two-tailed t-test). Scale bar: 25 μm. **(H)** Schematic illustration of the precocious SC disassembly observed in *Fbxo47* KO spermatocytes. The expression timing of H1t and H3S10P markers is shown.

Curiously, however, *Fbxo47* KO spermatocytes exhibited apparent diplotene-like chromosome morphology, despite the failure in reaching H1t positive mid pachytene (Fig. 5A). It is known that homolog synapsis is initiated at interstitial regions on the chromosome arm at zygotene, and that de-synapsis of homologs first starts at interstitial regions on the chromosome arm, while telomere regions are prone to be the last place of de-synapsis at diplotene (Bisig et al., 2012) (Qiao et al., 2012). This cytological difference readily distinguishes de-synapsed chromosomes at diplotene from un-synapsed ones at zygotene. Indeed, those *Fbxo47* KO spermatocytes with diplotene-like chromosome morphology apparently showed a typical feature of de-synapsis of homologs, wherein telomere regions retained homolog synapsis while interstitial regions were free from synapsis. To solve the paradox that *Fbxo47* KO spermatocytes showed diplotene-like chromosome morphology despite the failure of progressing beyond H1t-positive pachytene stage, we further analyzed the meiotic prophase population at P15 and P18 in the first wave of spermatogenesis of *Fbxo47* KO testes. Notably, diplotene-like cells (6.7 %) appeared in *Fbxo47* KO spermatocytes as early as P15, whereas the first wave of spermatogenesis was yet to pass beyond pachytene stage in the age-matched WT (Fig. 5C). HORMAD1 localizes along un-synapsed chromosomes before pachytene and de-synapsed chromosomes at diplotene, but dissociates from synapsed chromosomes (Shin et al., 2010) (Daniel et al., 2011) (Wojtasz et al., 2009). In *Fbxo47* KO spermatocytes, HORMAD1 dissociated from synapsed chromosomes at pachytene and re-localizes on de-synapsed chromosomes at diplotene-like stage as in those of WT (Fig. 5D), suggesting that localization of HORMAD1 on chromosomes was normally regulated. Histone H3 Ser10 phosphorylation (H3S10p) by Aurora B kinase of the chromosome passenger complex marks the centromeric region at diplotene and the whole chromosome at metaphase I (Parra et al., 2009) (Parra et al., 2003). In the control spermatocytes, the centromeric regions at diplotene were indicated by immunostaining of H3S10p (Fig. 5E). In contrast, H3S10p-positive centromeric regions were not observed in *Fbxo47* KO diplotene-like spermatocytes (Fig. 5E). This observation indicated that *Fbxo47* KO spermatocytes failed to reach *bona fide* diplotene stage of meiotic prophase, albeit exhibiting apparent homolog de-synapsis. Thus, we reasoned that even though homolog synapsis once occurred, it was destabilized during pachytene in *Fbxo47* KO spermatocytes. It should be mentioned that more zygotene and reciprocally less pachytene populations were observed in *Fbxo47* KO spermatocytes compared to WT at P15 and P18 (Fig. 5C). This implies that the process of homolog synapsis, at least in part, may be delayed in *Fbxo47* KO spermatocytes.

Mid-late pachytene spermatocytes acquire competency for meiotic prophase-Metaphase I transition indicted by the response to phosphatase inhibitor okadaic acid (OA) (Cobb *et al*., 1999). *In vitro* culture of isolated spermatocytes in the absence or presence of OA demonstrated that while the control spermatocytes progressed to diakinesis/metaphase I in the presence of OA, *Fbxo47* KO spermatocytes did not (Fig. 5F). Since *Fbxo47* KO spermatocytes were yet to acquire competency for OA-induced progression into metaphase I, even the most advanced *Fbxo47* KO spermatocytes remained in an earlier cell cycle stage compared to the control. These results suggested that the primary defect occured at zygotene or early pachytene stage in *Fbxo47* KO spermatocytes. Notably, TUNEL positive cells were observed in ∼21% of *Fbxo47* KO seminiferous tubules (Fig. 5G), suggesting that *Fbxo47* KO spermatocytes were consequently eliminated by apoptosis. Altogether, these results suggested that SC was prematurely disassembled in *Fbxo47* KO spermatocytes (Fig. 5H).

### *Fbxo47* KO spermatocytes show defects in meiotic recombination

Aforementioned results suggested that FBXO47 protein was required for stable maintenance of SC (Fig. 5). SC facilitates meiotic recombination that is executed by DSB formation and repair steps. Then SC is disassembled after the completion of crossover formation. Given that SC was prematurely destabilized in *Fbxo47* KO spermatocytes, we assumed two possibilities: (1) premature SC disassembly could be a result of early completion of meiotic recombination. (2) premature SC disassembly abolished the processes of meiotic recombination. To address these issues, we examined DSB formation and repair events by immunostaining of γH2AX. The first wave of γH2A is mediated by ATM after DSB formation at leptotene (Mahadevaiah et al., 2001), and disappears during DSB repair. The second wave of γH2A at zygotene is mediated by ATR that targets unsynapsed chromosomes (Royo et al., 2013). At zygotene, γH2AX signal appeared in *Fbxo47* KO spermatocytes in the same manner as WT (Fig. 6A), indicating that DSB formation normally occurred in *Fbxo47* KO spermatocytes. However, γH2AX signals largely persisted throughout the nuclei until pachytene-like and diplotene-like stages in *Fbxo47* KO spermatocytes, while they overall disappeared in WT pachytene spermatocytes except for retaining on the XY body (Fig. 6A). This observation suggested that DSB was still not repaired in *Fbxo47* KO diplotene-like spermatocytes. Furthermore, BRCA1, a marker of asynapsis (Scully et al., 1997) (Turner et al., 2004) (Broering et al., 2014), appeared along unsynapsed autosomal axes in zygotene *Fbxo47* KO spermatocytes as in those of WT (Fig. 6B). This suggests that meiotic silencing of unsynapsed chromatin (MUSC) was normally activated in *Fbxo47* KO spermatocytes. Crucially, in contrast to un-synapsed chromosomes in zygotene, BRCA1 was not observed along precociously de-synapsed chromosomes in *Fbxo47* KO diplotene-like spermatocytes (Fig. 6B). This suggests that MUSC was canceled in *Fbxo47* KO diplotene-like spermatocytes, presumably once homolog synapsis had successfully been achieved.

**Figure 6.**
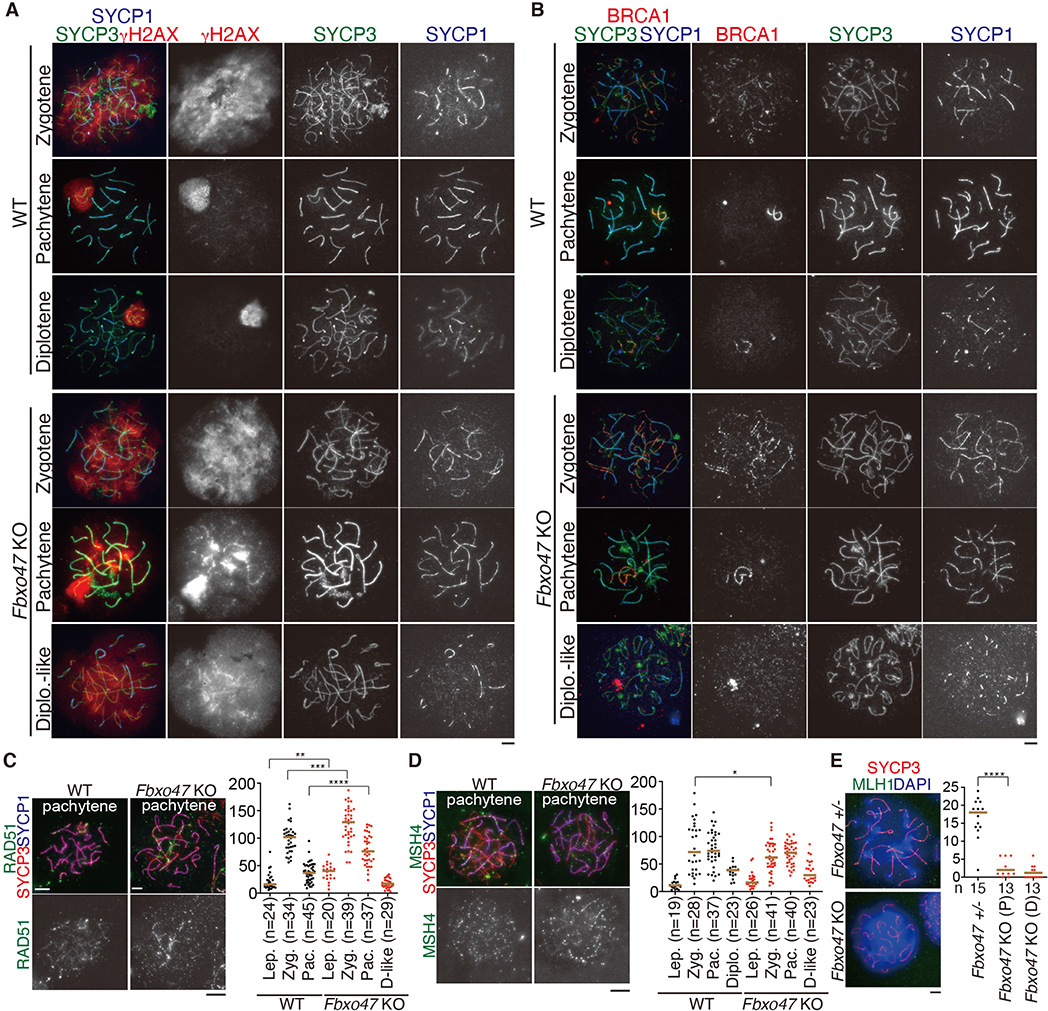
*Fbxo47* KO spermatocytes show defects in meiotic recombination. **(A)** Chromosome spreads of WT and *Fbxo47* KO spermatocytes were immunostained for SYCP3, SYCP1 and γH2AX. **(B)** Chromosome spreads of WT and *Fbxo47* KO spermatocytes were immunostained for SYCP3, SYCP1 and BRCA1. **(C)** Chromosome spreads of WT and *Fbxo47* KO spermatocytes were stained as indicated. Immunostained chromosome spread of pachytene spermatocytes are shown. The number of RAD51 foci is shown in the scatter plot with median (right). Statistical significance is shown by *p*-value (Mann-Whitney U-test). ****: p < 0.0001. ***: p < 0.001. **: p < 0.01. Lep.: leptotene, Zyg.: Zygotene, Pac.: Pachytene, Z-like: Zygotene-like, P-like: Pachytene-like, D-like: Diplotene-like. n: the number of cells examined. **(D)** Chromosome spreads of WT and *Fbxo47* KO spermatocytes were stained as indicated. The number of MSH4 foci is shown in the scatter plot with median (right). Statistical significance is shown by *p*-value (Mann-Whitney U-test). *: p < 0.05. **(E)** Chromosome spreads of *Fbxo47*+/- and *Fbxo47* KO spermatocytes were stained as indicated. The number of MLH1 foci is shown in the scatter plot with median (right). Statistical significance is shown by *p*-value (Mann-Whitney U-test). ***: p < 0.0001. Scale bars: 5 μm.

RAD51 facilitates the invasion of 3’-extended strand into the duplex of homolog at DSBs (Cloud et al., 2012) (Shinohara and Shinohara, 2004). In accordance with the persistent DSBs in *Fbxo47* KO (Fig. 6A), the number of RAD51 foci was significantly increased in *Fbxo47* KO spermatocytes (Fig. 6C). Reciprocally, the number of MSH4 foci was decreased in *Fbxo47* KO spermatocytes (Fig. 6D). These observations suggest that although RAD51 was normally loaded onto DSBs, the processes of homologous recombination-mediated repair were delayed or blocked in the absence of FBXO47. Accordingly, the number of MLH1 foci, a marker of crossover (CO), was significantly reduced in *Fbxo47* KO pachytene-like spermatocytes compared to WT pachytene spermatocytes (Fig.6E). This implies that crossover recombination was incomplete in the absence of FBXO47. Altogether, precocious disassembly of SC was a cause of the defect in meiotic recombination rather than a result of early completion of meiotic recombination.

## Discussion

### FBXO47 stabilizes homolog synapsis independently of SCF in mouse

We have shown that FBXO47 is required for the maintenance of homolog synapsis during prolonged meiotic prophase. *Fbxo47* KO spermatocytes showed precocious de-synapsis, albeit exhibiting apparently “diplotene-like” morphology (Fig. 5B). Although this phenomenon in *Fbxo47* KO spermatocytes was partly similar to that observed in conditional *Skp1* KO (Guan *et al*., 2020), marked phenotypic differences were observed between *Fbxo47* KO and *Skp1* KO spermatocytes. In *Skp1* KO testis, late pachytene spermatocytes are absent and concurrently diplotene spermatocytes are increased. *Skp1* KO spermatocytes at least reach H1t positive mid-pachytene in terms of cell cycle, but most of them contain de-synapsed chromosomes at pericentric end termed “Y pachynema”. Thus, *Skp1* KO spermatocytes show precocious de-synapsis and pachyetene exit. In contrast, *Fbxo47* KO spermatocytes failed to reach H1t positive mid-pachytene (Fig. 5A). Although apparent diplotene-like morphology of homolog chromosomes appeared in *Fbxo47* KO spermatocytes (Fig. 5B), “Y pachynema” was not observed in *Fbxo47* KO, unlike in *Skp1* KO spermatocytes. Thus, *Fbxo47* KO spermatocytes show precocious desynapsis despite the failure of progression beyond pachyetene. These results suggested that primary defect in *Fbxo47* KO spermatocyte occurred at earlier cell cycle stage than *Skp1* KO spermatocytes. HORMAD1 localizes along unsynapsed and de-synapsed chromosomes during meiotic prophase (Shin *et al*., 2010) (Daniel *et al*., 2011), and dissociates from synapsed chromosomes by the action of TRIP13 AAA ATPase (Wojtasz *et al*., 2009). Wheas HORMAD1 persists both in synapsed and desynapsed chromosomes in *Skp1* KO spermatocyte, localization of HORMAD1 on chromosomes was normally regulated in *Fbxo47* KO (Fig. 5D). Thus, precocious desynapsis could be derived at least in part from failure of HORMAD1 removal in *Skp1* KO and from different mechanism in *Fbxo47* KO. Moreover, while DSB repair process indicated by γH2AX staining (Fig. 6A) was impaired both in *Fbxo47* KO and in *Skp1* KO spermatocytes, the extent of crossover formation was different between them. Whereas significant number of MLH1 foci were observed in mid-late pachytene and diplotene spermatocytes in *Skp1* KO, MLH1 foci were rarely observed in pachytene and diplotene-like spermatocytes in *Fbxo47* KO (Fig. 6E). Thus, meiotic recombination and crossover formation were more progressed in *Skp1* KO than in *Fbxo47* KO.

Fbox-domain containing proteins confers substrate specificity to SCF E3 ubiquitin ligase (Jin *et al*., 2004) (Reitsma *et al*., 2017). Altough FBXO47 possesses a putative Fbox domain, it was not detected in the SKP1 immunoprecipitates from *Skp1-3xFLAG-HA* KI testes (Fig. 2D, Supplementary Data1). Reciplocally, SKP1 was not detected in the immunoprecipitates from *Fbxo47 -3xFLAG-HA* KI testes (Fig. 1H, Fig S2). Furthermore, while SKP1 localized along lateral element (LE) of synapsed chromosomes (Fig. 2B) (Guan *et al*., 2020), FBXO47 protein did not show such a specific localization pattern on the chromosome (Fig. S3B). Although we cannot formally exclude a possibility that FBXO47 is incorporated as a substrate recognition subunit in SCF under specific regulation, our results suggest that FBXO47 may not be incorporated in the function of SCF, and rarther FBXO47 may function independently of SCF in spermatocytes.

### Distinct functions of FBXO47 homologs in diverse organisms

Fbxo47 homologues and other distant F-box proteins have been implicated in meiotic prophase progression in various species. Although defects accompanying DSB repair and crossover are similarly observed in mouse and *C. elegans Fbxo47* mutants, the primary causes are assumed to be different. In *C. elegans,* PROM-1 encodes *Fbxo47* homolog. In *C. elegans*, organization of gonadal germline is devided into mitotic/meiotic entry zone, transition zone corresponding to zyotene, and pachytene zone. *Prom-1* mutant showed delayed and asynchronous initiation of homolog pairing, so that distinct transition zone was missing and meiotic entry zone was rather extended (Jantsch *et al*., 2007) with attenuating CHK-2 activity (Mohammad et al., 2018) (Baudrimont et al., 2021). Further, PROM-1 was proposed to down regulate mitotic cell cycle proteins such as Cyclin E homolog CYE-1 at meiotic entry, independently of promoting homolog pairing as a positive regulator of CHK-2 kinase (Mohammad *et al*., 2018).Thus, PROM-1 functions very early in meiotic prophase in *C. elegans*, which is similar to our observation in mice (Fig. 5). In *prom-1* meiocytes however, homolog pairing was defective and non-homologous synapsis was consequently pronounced in autosomes but not in X chromosome. Thus, PROM-1 is implicated in promoting autosome homolog pairing. This is a contrast to our observation, in which homolog synapsis once took place normally, followed by premature desynapsis in *Fbxo47* KO sepermatocytes (Fig. 5B).

In the teleost fish medaka, *fbxo47* mutant XX germ cells exhibit abnormally condensed chromosomes in ovaries and fail to undergo oogenesis after diplotene, showing that the sexual fate of XX germ cells turns into spermatogenesis (Kikuchi *et al*., 2020). Thus, *fbxo47* is involved in the regulation of cell division in ovaires, and in turn the suppression of spermatogenesis in female germ cells in medaka. The germline feminization under *fbxo47* is mediated at least by two downstream transcription factors *lhx8b* and *figla* during early meiotic prophase in medaka. Despite the phenotypical similarities and differences observed in the mutants of Fbxo47 homologs in diverse organisms, FBXO47 homologs commonly act during meiotic prophase, although at different time points.

### Distinct interpretations on the function of FBXO47 in mouse

Previous study showed that mouse FBXO47 interacts with SKP1 and telomere binding proteins, TRF1 and TRF2 (Hua *et al*., 2019). According to the study, FBXO47 was localized to telomeres during meiotic prophase. Furthermore, TRF2 were destabilized and telomeres were detached from the nulclear envelope in *Fbxo47* KO spermatocytes, causing defects in telomere bouquet formation (Hua *et al*., 2019). Those observations led to propose that FBXO47 binds to telomeric proteins TRF1 and TRF2, and plays a role in protecting TRF2 from destruction (Hua *et al*., 2019). However, we failed to detect either TRF1 or TRF2 in FBXO47 immunoprecipitates from testis chromatin fraction (Fig. S3A). Further, we failed to observe localization of FBXO47 to telomeres (Fig. S3B) and detachment of telomeres from nuclear envelope in *Fbxo47* KO spermatocytes (Fig. S3C), which contrast to the previous report (Hua *et al*., 2019). Since the frequency of bouquet formation was quite low even in WT spermatocytes in mouse (Fig. S3D), as shown in our previous study (Ishiguro et al., 2014), the potential defect in bouquet formation in *Fbxo47* KO spermatocytes further needs to be evaluated. Furthermore, our SKP1 immunoprecipitation from *Skp1-3xFLAG-HA* KI testes (Fig. 2D, Supplementary Data1) and reciplocal FBXO47 immunoprecipitation from *Fbxo47-3xFLAG-HA* KI testes (Fig. 1H, Fig S2) failed to show supporting evidence that FBXO47 serves as a subunit of SCF. Although we do not know the exact reason for the discrepancies between the two studies with similar histogical phenotype of the seminiferous tubules in *Fbxo47* KO testes, subtle differences in the dection and assay conditions or mice that were used could account for the differences in the observations.

### Distant F-box proteins are involved in homolog synapsis

SCF and F-box proteins are involved in the process of homolog synapsis during meiotic prophase in diverse organisms. In plants, although no *Fbxo47* homolog exsit, distant F-box proteins are involved in homolog synapsis. In rice plant (*Oryza sativa*), MEIOTIC F-BOX (MOF) encodes a F-BOX protein, and interacts with OSK1, a homolog of SKP1 (He *et al*., 2016). MOF acts as a subunit of SCF and localizes on the chromosome during meiotic prophase. In *mof* mutant male meiocytes, telomeres were not clustred and homolog synapsis was lost as indicated by complete absence of ZEP1, a transverse filament of SC. Thus, MOF plays a role in telomere bouquet formation during homolog pairing in male meiocyte. In rice plant, ZYGOTENE1 (ZYGO1) encodes another F-box protein that has a limited similarity to mouse FBL12 (Zhang *et al*., 2017). In *zygo1* mutant, polarized enrichment of OsSAD1, a SUN-domain containing protein, along nuclear envelope was lost and full-length homolog pairing was consequently impaired. This led to defective DSB repair of meiotic recombination, causing both male and female sterility in *zygo1* mutant. Thus, ZYGO1 also plays a role in telomere bouquet formation during homolog pairing in rice plant. These studies suggest that rice F-box proteins MOF and ZYGO1 act as a SCF component, and play a role in bouquet formation rather than in the process of SC formation, which is different to the role of mouse FBXO47 in SC maintenance.

In budding yeast, a F-box protein Cdc4 acts as a substrate subunit of SCF durimg meiotic prophase. SCF ^Cdc4^ is assumed to regulate SC assembly by counteracting the Pch2 (TRIP13 in mammals)-dependent negative action that induces SC disassembly (Zhu *et al*., 2021). It is proposed that SCF ^Cdc4^ targets the putative negative regulator of SC assembly toward degradation, and in turn stabilizes SC. Although how Pch2 itself or its downstream factors is counteracted by SCF ^CDC4^ remines elusive, F-box protein Cdc4 acts for the maintenance of SC in budding yeast.

In *Drosophila* female, knockdown of *SkpA,* a *Skp1* homolog, caused premature disassembly of SC (Barbosa *et al*., 2021). Depletion of F-box proteins, Fbxo42 and Slmb/βTrcp, showed imcomplete formation and precousious disassembly of SC, which was similar to the observation in *Fbxo47* KO mouse. PP2A catalytic (C) subunit and structural (A) subunit were identified as a candidate substrate of Fbxo42. Since overexpression of a PP2A subunit Wrd (B56) phenocopied Fbxo42 knockdown, the SCF ^Fbxo42^ is assumed to satabilize SC by restricting PP2A-Wrd (B56) association. In these regards, *Drosophila* Fbxo42 and budding yeast Cdc4 share a similar role to mouse FBXO47 in maintaining SC stability.

Previous studies showed PLK1 mediated-phosphorylation regulate SC disassembly in mouse (Jordan et al., 2012), and PP2A phosphatase inihibitor OA promotes premature exit from pachytene and SC disassembly (Cobb *et al*., 1999). Thus, phosphorylation level of SC regulates its stability during meiotic cell cycle. Given that FBXO47 exists in the cytosol rather than localizing to the chromatin (Fig. 1H), it is possibile that FBXO47 may protect the SC directly or indirectly from a putative destabilizer that regulates the phosphorylation level of SC during early meiotic prophase (Fig. 7). It is still a large enigma how FBXO47 acts for preventing premature SC disassembly, and further investigation is required for understanding the precise mechganism of FBXO47 function.

**Figure 7.**
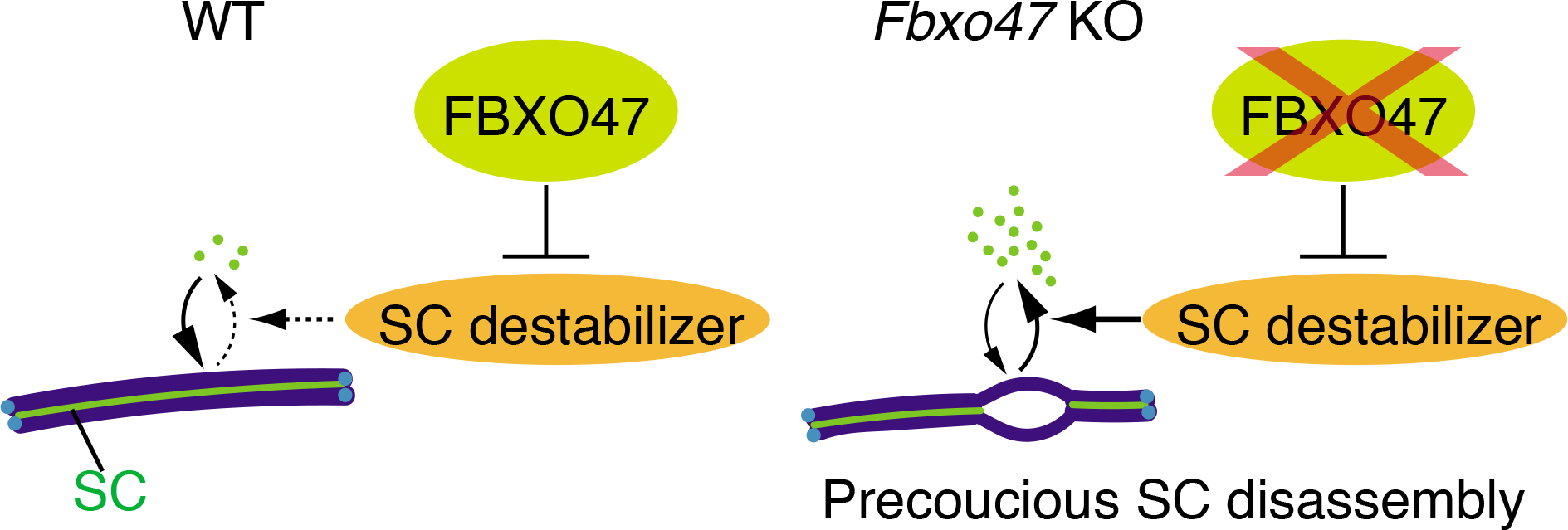
A model of FBXO47 function to prevent premature SC disassembly. Schematic illustration how FBXO47 may protect SC from a putative destabilizer during early meiotic prophase.

## Acknowledgments

The authors thank Kaho Okamura (Kumamoto University) for technical support, and Marry Ann Handel for provision of H1t antibody. This work was supported in part by KAKENHI grant (#21K15018 to N.T.), KAKENHI grant (#19K06642 to Y.T.), KAKENHI grant (#20K22638 to R.S.), and KAKENHI grants (#19H05743, #20H03265, #20K21504, #JP 16H06276 to K.I.) from MEXT Japan; Grant from AMED PRIME (21gm6310021h0001 to K.I.). Grants from The Sumitomo Foundation; The Naito Foundation, Astellas Foundation for Research on Metabolic Disorders; Daiichi Sankyo Foundation of Life Science; The Uehara Memorial Foundation; The NOVARTIS Foundation (Japan) for the promotion of Science; Takeda Science Foundation (to K.I.).

## Author contributions

N.Tanno, K.T. performed the cytological and biochemical analyses. R.S. performed reanalysis of scRNA-seq data. N.Tani performed MS analuses. Y.T.H. performed the RT-PCR. K.A. designed the knockout mice. S.F. performed histological analyses. N.Takeda assisted oocyte experiments. K.I. supervised experiments, conducted the study and wrote the manuscript.

## Declaration of interests

The authors declare no competing interests.

**Supplementary Data1 (Excel). Whole list of identified proteins by MS analyses of FBXO47 IP and SKP1 IP from testis extracts**

Coloidal blue stained gel after running the samples is shown in the first tab. Gel was cut into pieces before LC-MS/MS analysis. Full list of proteins identified by the LC-MS/MS analysis are shown in the second tab. The proteins are listed with UniProt accession number, the number of peptide hits and Mascot scores. In the third tab, proteins are presented after excluding the proteins detected in the control mock purification, IgG and keratin.

**Supplementary Data2 (Excel). The source data for statistics**

The source data (for Fig.1B, Fig.1F, Fig.4C, Fig.4G, Fig. 4I, Fig. 4J, Fig. 5A, Fig. 5C, Fig. 5F, Fig.5G, Fig. 6C, Fig. 6D, Fig. 6E) are shown in the tabs.

## STAR Methods

### Lead Contact and Material Availability

Further information and requests for the resources and reagents should be directed to and will be fulfilled by the Lead Contact, Kei-ichiro Ishiguro.

All data supporting the conclusions are present in the paper and the supplementary materials. The source data (for Fig.1B, Fig.1F, Fig.4C, Fig.4G, Fig. 4I, Fig. 4J, Fig. 5A, Fig. 5C, Fig. 5F, Fig.5G, Fig. 6C, Fig. 6D, Fig. 6E, FigS3D) are provided in Supplementary data2 (Excel). The original images for all of the figures in this paper are deposited in public depository.

The ChIP-seq data of MEIOSIN and STRA8 are described in our previous study (Ishiguro *et al*., 2020) and available in the DDBJ Sequence Read Archive (DRA) under accession number DRA007066, DRA007778, DRA009056. Mouse lines generated in this study have been deposited to Center for Animal Resources and Development (CARD), *Fbxo47* Ex3-11Δ knockout mouse (ID 2777), *Fbxo47-3xFLAG-HA* knock-in mouse (ID 2972), and *Skp1-3xFLAG-HA* knock-in mouse (ID 2638). Plasmid expression vectors generated in this study have been deposited to RIKEN BRC: pET28c-*Fbxo47*-C (aa272-451) (ID RDB192639) and pET28c-*Fbxo47*-M (aa174 -316) (ID RDB19264). The antibodies are available upon request. There are restrictions to the availability of antibodies due to the lack of an external centralized repository for their distribution and our need to maintain the stock. We are glad to share antibodies with reasonable compensation by the requestor for its processing and shipping. All unique/stable reagents generated in this study are available from the Lead Contact with a completed Materials Transfer Agreement.

### Experimental Model and Subject Details

#### Animals

*Fbxo47* Ex3-11Δ knockout and *Fbxo47-3xFLAG-HA* knock-in mice were C57BL/6 background. *Skp1-3xFLAG-HA* knock-in mouse was congenic with C57BL/6 background. Male mice were used for immunoprecipitation of testis extracts, histological analysis of testes, immunostaining of testes, and RT-PCR experiments. Female mice were used for histological analysis of the ovaries, and immunostaining experiments. Whenever possible, each knockout animal was compared to littermates or age-matched non-littermates from the same colony, unless otherwise described. Animal experiments were approved by the Institutional Animal Care and Use Committee (approval F28-078, A30-001, A28-026, A2020-006).

### Method Details

#### Generation of *Fbxo47* knockout mice and genotyping

*Fbxo47* knockout mouse was generated by introducing Cas9 protein (317-08441; NIPPON GENE, Toyama, Japan), tracrRNA (GE-002; FASMAC, Kanagawa, Japan), synthetic crRNA (FASMAC), and ssODN into C57BL/6N fertilized eggs using electroporation. For generating *Fbxo47* Exon3-11 deletion (Ex3-11Δ) allele, the synthetic crRNAs were designed to direct TACACCTAGTGATAGCACTT(GGG) of the *Fbxo47* intron 2 and AGAGCACTAGTCACTGAATG(CGG) in the 3’-neighboring region of the Exon11. ssODN: 5’- GCTCAAAGTAAGCAAAGCAACAGAGAGCACTAGTCACTGATTATTTATTCACTTGGGATGCTGAGGAGGCAAATTGCCAGGTGTTTGAAGC

-3’ was used as a homologous recombination template.

The electroporation solutions contained (10μM of tracrRNA, of synthetic crRNA, 0.1μg/μl of Cas9 protein, 1μg/μl of ssODN) for *Fbxo47* knockout in Opti-MEM I Reduced Serum Medium (31985062; Thermo Fisher Scientific). Electroporation was carried out using the Super Electroporator NEPA 21 (NEPA GENE, Chiba, Japan) on Glass Microslides with round wire electrodes, 1.0 mm gap (45-0104; BTX, Holliston, MA). Four steps of square pulses were applied (1, three times of 3 mS poring pulses with 97 mS intervals at 30 V; 2, three times of 3 mS polarity-changed poring pulses with 97 mS intervals at 30 V; 3, five times of 50 mS transfer pulses with 50 mS intervals at 4 V with 40% decay of voltage per each pulse; 4, five times of 50 mS polarity-changed transfer pulses with 50 mS intervals at 4 V with 40% decay of voltage per each pulse).

The targeted *Fbxo47* Ex3-11Δ allele in F0 mice were identified by PCR using the following primers:

Fbxo47-F1: 5’-TCCTCTCTCTGTCTCTTTATTCAACAG-3’ and Fbxo47-R1: 5’-TGCTAAGAAGGTGGTAAAGAATGTGAC-3’ for the knockout allele (825 bp). Fbxo47-F3: 5’- TCTGACCATGAACGCTATCTCTTCC-3’ and Fbxo47-R1 for wild-type allele (503 bp). The PCR amplicons were verified by sequencing. Primer sequences are listed in Table S1.

#### Generation of *Fbxo47-3xFLAG-HA* knock-in mice and genotyping

*Fbxo47-3xFLAG-HA* knock-in mouse was generated by introducing Cas9 protein, tracrRNA, synthetic crRNA, and ssODN into C57BL/6N fertilized eggs using electroporation as described above. The synthetic crRNA was designed to direct ACGCTATCTCTTCCTAAGTC(AGG) of the *Fbxo47*.

ssODN:

5’-GAACTTCCATAAGGAGGTGCTGTATCTGACCATGAACGCTATCTCTTCCGGAGACTACAAAGACCATGACGGTGATTATAAAGATCATGACATCGATTACAAGGATGACGATGACAAGGGATACCCCTACGACGTGCCCGACTACGCCTAAGTCAGGAAGCTTGTGTCCCTCTGGACTGGCATTCAGGGGAGTGATGCC-3’

was used as a homologous recombination template.

The targeted *Fbxo47-3xFLAG-HA* knock-in allele in F0 mice were identified by PCR using the following primers:

Fbxo47-F4: 5’-TCTGTTCCATCTTCTCCATGCTCAGGC-3’ and Fbxo47-R3: 5’-TGAAGAGCCAGAACTTGTTTTCCAG-3’ for the knock-in allele (396 bp), and for wild-type allele (294 bp). The PCR amplicons were verified by sequencing. Primer sequences are listed in Table S1.

#### Generation of *Skp1-3xFLAG-HA* knock-in mouse and genotyping

The targeting vector was designed to insert 3xFLAG-HA-3’UTR in frame with the coding sequence into the Exon 6 of the *Skp1* genomic locus. Targeting arms of 1225bp and 1481bp fragments, 5’ and 3’ of the Exon 6 of *Skp1* gene respectively, were generated by PCR from mouse C57BL/6 genomic DNA and directionally cloned flanking p*GK-Neo*-polyA and *DT-A* cassettes. The 5’ arm was followed by nucleotide sequences encoding 3xFLAG, HA and the 3’UTR of *Skp1* gene. TT2 ES cells were co-transfected with the targeting vector and pX330 plasmids (Addgene) expressing Crispr-gRNAs directing GCTGGCATTGACTCGGGGTA(ggg) and CGCCACCATACCCGGTGATT (tgg), which locate at the 3’ region of the Exon 6 of *Skp1* gene. The G418-resistant ES clones were screened for homologous recombination with the *Skp1* locus by PCR using primers SKP1_5Arm_F2: 5’-

GGTCAGCAACACTGCTGAACAGCTTG-3’ and

KI96ES-19814R-HA: 5’- GGGCACGTCGTAGGGGTATCCCTTG -3’ for the left arm (1909 bp); pKO2-3armF: 5’-AGGAACTTCGGAATAGGAAC-3’ and

SKP1_RightArm_R2: 5’-TGCAGTGGAGGCTCAGTCCAGCTTC-3’ for the right arm (1897 bp).

The homologous recombinant cells were isolated and chimeric mice were generated by aggregation (host ICR) of recombinant ES cells. Chimeric males were mated to C57BL/6N females and the progenies were genotyped by PCR using the primers:

SKP1onL2_F2: 5’- ATCATTGTTCCCAGGTGGAG -3’ and SKP1onRight_R1: 5’- GACTAGAACAAGATGACAGG -3’

for the knock-in allele (2078 bp) and the WT allele (1275bp). Primer sequences are listed in Table S1.

#### Histological Analysis

Testes, caudal epididymis and ovaries were fixed in Bouin’s solution, and embedded in paraffin. Sections were prepared on CREST-coated slides (Matsunami) at 6 μm thickness. The slides were deparaffinized and stained with hematoxylin and eosin.

For Immunofluorescence staining, testes were embedded in Tissue-Tek O.C.T. compound (Sakura Finetek) and frozen. Cryosections were prepared on the CREST-coated slides (Matsunami) at 8 μm thickness, and then air-dried. The serial sections of frozen testes were fixed in 4% paraformaldehyde in PBS for 5 min at room temperature and washed briefly in PBS. After washing, the serial sections were permeabilized in 0.1% TritonX100 in PBS for 5 min.

The sections were blocked in 3% BSA/PBS, and incubated at room temperature with the primary antibodies in a blocking solution. After three washes in PBS, the sections were incubated for 1 h at room temperature with Alexa-dye-conjugated secondary antibodies (1:1000; Invitrogen) in a blocking solution. PNA lectin staining was done using FITC-conjugated Lectin from *Arachis hypogaea* (IF, 1:1000, Sigma: L7381). TUNEL assay was performed using MEBSTAIN Apoptosis TUNEL Kit Direct (MBL 8445). DNA was counterstained with Vectashield mounting medium containing DAPI (Vector Laboratory).

#### Immunostaining of spermatocytes

Surface-spread nuclei from spermatocytes were prepared by the dry down method as described (Peters et al., 1997) (Takemoto et al., 2020) with modification. The slides were then air-dried and washed with water containing 0.1 % TritonX100 or frozen for longer storage at -30°C. The slides were permeabilized in 0.1% TritonX100 in PBS for 5 min, blocked in 3% BSA/PBS, and incubated at room temperature with the primary antibodies in 3% BSA/PBS. After three washes in PBS, the sections were incubated for 1 h at room temperature with Alexa-dye-conjugated secondary antibodies (1:1000; Invitrogen) in a blocking solution. For bouquet counting, cells were suspended in PBS without hypotonic treatment and structurally preserved nuclei of spermatocytes were prepared by cytospin at 1000rpm for 5min (Thermofisher). Cells were fixed with 4% PFA in PBS for 5 min. The slide grasses were washed with PBS containing 0.1% Triton-X100 in PBS. After washing with PBS, immunofluorescence staing was perfomed immediately. DNA was counterstained with Vectashield mounting medium containing DAPI (Vector Laboratory).

#### Imaging

Immunostaining images were captured with DeltaVision (GE Healthcare). The projection of the images was processed with the SoftWorx software program version 7.2.1 (GE Healthcare). All images shown were Z-stacked. Bright field images and immunofluorescent images for counting seminiferous tubules, were captured with BIOREVO BZ-X710 (KEYENCE), and processed with BZ-H3A program. XY-stitching capture by 10x objective lens was performed for multiple-point color images using BZ-X Wide Image Viewer. Images were merged over the field using BZ-H3A Analyzer (KEYENCE). If the SYCP3 image was too dim for counting the SYCP3+ seminiferous tubules, the contrast of the color channel used for SYCP3 was enhanced in the XY-stitched image.

#### *In vitro* oocyte culture and Giemsa staining of metaphase chromosome spread

Ovaries collected from 4-week-old female mice were used after 46 to 48 h of treatment with 5 IU of pregnant mare serum gonadotropin. GV oocytes were isolated by puncturing the follicles in M2 medium (Sigma MR-015). The GV oocytes were cultured in M16 medium (Sigma MR-016) in a 5% CO_2_ atmosphere at 37°C for 6hours. For Giemsa staining of metaphase chromosome spread, oocytes were exposed to 0.5% Pronase (MERCK 10165921001) to remove the zona pellucida, and treated in hypotonic buffer containing 1% sodium citrate/0.1% PVA for 15min. The oocytes and oocyte-like cells were placed on the slides, fixed in the Carnoy’s Fixative (75 % Methanol, 25% Acetic Acid), and stained in 3% Giemsa solution for 30min.

#### Culture of OA-induced Meta I spermatocyte

Culture of OA-induced Meta I spermatocytes were performed as described (Wiltshire et al., 1995). The isolated spermatocytes were cultured in the presence or absence of 5 μM okadaic acid (OA) for 3 h.

#### Antibodies

The following antibodies were used for immunoblot (IB) and immunofluorescence (IF) studies: mouse anti-FLAG M2 (Sigma-Aldrich F1804), rabbit anti-HA (IB, IF, 1:1000, Abcam: ab9110), rabbit anti-Actin (IB, 1:1000, Sigma-Aldrich A2066), mouse anti-MLH1 (IF, 1:500, BD Biosciences: 551092), rabbit anti-H3S10p (IF, 1:2000, Abcam ab5176), rabbit anti-Histone H3 (IB, 1:1000, Abcam ab1791), rabbit anti-SYCP1 (IF, 1:1000, Abcam ab15090), mouse anti-gH2AX (IF, 1:1000, Abcam ab26350), rabbit anti-RAD51 (IF, 1:500, Santa Cruz: SC-8349), rabbit anti-MSH4 (IF, 1:500, Abcam ab58666), rabbit anti-SKP1 (IB, 1:1000, Abcam ab10546), rabbit anti-HORMAD1 (IF, 1:1000, ProteinTech 13917-1-AP), goat anti-Lamin B (IF, 1:1000, Santa Cruz: SC-6216), mouse anti-TRF1(Shibuya et al., 2014) (IF, 1:1000), rabbit anti-TRF1 (Shibuya *et al*., 2014) (IB, 1:1000), rabbit anti-TRF2 (IB, 1:1000, NB110-57130), mouse anti-SYCP1 (IF, 1:1000) (Ishiguro et al., 2011), rat anti-SYCP3 (Ishiguro *et al*., 2020) (IF, 1:1000), gunia pig anti-SYCP3 (Ishiguro *et al*., 2020) (IF, 1:2000), rabbit anti-BRCA1 (IF, 1:500, kindly provided by Satoshi Namekawa), guinea pig anti-H1t (IF, 1:2000, kindly provided by Marry Ann Handel).

#### Production of antibodies against FBXO47

Polyclonal antibodies against mouse FBXO47 C-terminal (aa272-451) were generated by immunizing rabbits and a guinea pig. FBXO47 middle region (aa174-316) were generated by immunizing a rabbit. His-tagged recombinant proteins of FBXO47 middle region (aa174-316) and C-terminal (aa272-451) were produced by inserting cDNA fragments in-frame with pET19b and pET28c (Novagen) respectively in *E. coli* strain BL21-CodonPlus (DE3)-RIPL (Agilent), solubilized in a denaturing buffer (6 M HCl-Guanidine, 20 mM Tris-HCl pH 7.5) and purified by Ni-NTA (QIAGEN) under denaturing conditions. The antibodies were affinity-purified from the immunized serum with immobilized antigen peptides on CNBr-activated Sepharose (GE healthcare).

#### PCR with reverse transcription

Total RNA was isolated from tissues and embryonic gonads using TRIzol (Thermo Fisher). cDNA was generated from total RNA using Superscript III (Thermo Fisher) followed by PCR amplification using Ex-Taq polymerase (Takara) and template cDNA.

For RT-qPCR, total RNA was isolated from WT (n = 3) and *Meiosin* KO (n = 3) testes, and cDNA was generated as described previously (Ishiguro *et al*., 2020). *Fbxo47* cDNA was quantified by ⊗CT method using TB Green Premix Ex Taq II (Tli RNaseH Plus) and Thermal cycler Dice (Takara), and normalized by *GAPDH* expression level.

qPCR was performed in duplicates, and the average ddCt value was calculated for each cDNA sample. The expression level of *Fbxo47* was divided by that of *GAPDH* to give the relative expression level of *Fbxo47* to *GAPDH.* Relative expression level of *Fbxo47* to *GAPDH* was normalized to 1 for a given P10 WT sample.

Sequences of primers used for RT-PCR were as follows: GAPDH-F: 5’-TTCACCACCATGGAGAAGGC-3’ GAPDH-R: 5’-GGCATGGACTGTGGTCATGA-3’ Gapdh_F2: 5’-ACCACAGTCCATGCCATCAC-3’ Gapdh_R2: 5’-TCCACCACCCTGTTGCTGTA-3’ Gapdh_Ex6F: 5’-GGTTGTCTCCTGCGACTTCA-3’

Gapdh_mRNAR: 5’-GCCGTATTCATTGTCATACCAGG-3’ *Fbxo47*-F 1443F: 5’-GCATAGCAAATGCTTTTGCCTGTG-3’ *Fbxo47*-R 1605R: 5’-GAGATAGCGTTCATGGTCAGATAC-3’

Primer sequences are listed in Table S1.

#### Preparation of testis extracts and immunoprecipitation

Testis chromatin-bound and -unbound extracts were prepared as described previously (Ishiguro *et al*., 2014). Briefly, testicular cells were suspended in low salt extraction buffer (20 mM Tris-HCl pH 7.5, 100 mM KCl, 0.4 mM EDTA, 0.1% TritonX100, 10% glycerol, 1 mM β-mercaptoethanol) supplemented with Complete Protease Inhibitor (Roche). After homogenization, the soluble chromatin-unbound fraction was separated after centrifugation at 100,000*g* for 10 min at 4°C. The chromatin bound fraction was extracted from the insoluble pellet by high salt extraction buffer (20 mM HEPES-KOH pH 7.0, 400 mM KCl, 5 mM MgCl_2_, 0.1% Tween20, 10% glycerol, 1 mM β-mercaptoethanol) supplemented with Complete Protease Inhibitor. The solubilized chromatin fraction was collected after centrifugation at 100,000*g* for 10 min at 4°C.

#### Immuno-affinity purification

Immuno-affinity purification was performed with anti-FLAG M2 monoclonal antibody-coupled magnetic beads (Sigma-Aldrich M8823) from the testis chromatin-bound and -unbound fractions of *Fbxo47*-*3xFLAG-HA* knock-in mice and *Skp1*-*3xFLAG-HA* knock-in mice (14 to 21-day old). For negative control, mock immuno-affinity purification was done from the testis chromatin-bound and -unbound fractions from the age-matched wild type mice. The beads were washed with high salt extraction buffer for chromatin-bound proteins and low salt extraction buffer for chromatin-unbound proteins. The anti-FLAG-bound proteins were eluted by 3xFLAG peptide (Sigma-Aldrich). The second immuno-affinity purification was performed anti-HA 5D8 monoclonal antibody-coupled Magnet agarose (MBL M132-10). The bead-bound proteins were eluted with 40 µ l of elution buffer (100 mM Glycine-HCl pH 2.5, 150 mM NaCl), and then neutralized with 4 µ l of 1 M Tris-HCl pH 8.0.

The immunoprecipitated proteins were run on 4-12 % NuPAGE (Thermo-Fisher) in MOPS-SDS buffer and silver-stained with Silver Quest (Thermo-Fisher), immunoblotted or analyzed by LC-MS/MS. For the immunoblot of whole testes extracts from WT, *Fbxo47* KO, and *Fbxo47-3FH* KI mice, lysates were prepared in RIPA buffer and run on 8% Laemmli SDS-PAGE in Tris-Glycine-SDS buffer. Immunoblot images were developed using ECL prime (GE healthcare) and captured by FUSION Solo (VILBER).

#### Mass spectrometry

The immunoprecipitated proteins were run on 4-12 % NuPAGE (Thermo Fisher) by 1 cm from the well and stained with SimplyBlue (Thermo Fisher) for in-gel digestion. The gel containing proteins was excised, cut into approximately 1mm sized pieces. Proteins in the gel pieces were reduced with DTT (Thermo Fisher), alkylated with iodoacetamide (Thermo Fisher), and digested with trypsin and Lysyl endopeptidase (Promega) in a buffer containing 40 mM ammonium bicarbonate, pH 8.0, overnight at 37°C. The resultant peptides were analyzed on an Advance UHPLC system (ABRME1ichrom Bioscience) connected to a Q Exactive mass spectrometer (Thermo Fisher) processing the raw mass spectrum using Xcalibur (Thermo Fisher Scientific). The raw LC-MS/MS data was analyzed against the NCBI non-redundant protein/translated nucleotide database restricted to *Mus musculus* using Proteome Discoverer version 1.4 (Thermo Fisher) with the Mascot search engine version 2.5 (Matrix Science). A decoy database comprised of either randomized or reversed sequences in the target database was used for false discovery rate (FDR) estimation, and Percolator algorithm was used to evaluate false positives. Search results were filtered against 1% global FDR for high confidence level. All full lists of LC-MS/MS data are shown in Supplementary Data1 (Excel file).

#### ChIP-seq Data and Public RNA-seq data Analysis

MEIOSIN ChIP-seq data described in our previous study (Ishiguro *et al*., 2020) was analyzed for the *Fbxo47* locus. MEIOSIN binding site was shown along with genomic loci from Ensembl on the genome browser IGV.

#### Single cell RNA-seq Data Analysis

The scRNA-seq data of fetal ovaries was derived from DRA 011172 (Shimada *et al*., 2021). 10xGenomics Drop-seq data of mouse adult testis was derived from GEO: GSE109033 (Hermann *et al*., 2018). Reanalyses of scRNA-seq data were conducted using the Seurat package for R (v.3.1.3) (Stuart et al., 2019) and pseudotime analayses were conducted using monocle package for R: R (ver. 3.6.2), RStudio (ver.1.2.1335), and monocle (ver. 2.14.0) (Qiu et al., 2017) following developer’s tutorial.

#### Quantification and Statistical analysis

Statistical analyses, and production of graphs and plots were done using GraphPad Prism8 (version 8.4.3) or Microsoft Excel (version 16.48).

---

Figure 1B Testis RNA was obtained from P8 WT (3 animals), P10 WT (3 animals), *Meiosin* KO (3 animals). qPCR was performed in duplicates, and the average ddCt value was calculated for each cDNA sample. The expression level of *Fbxo47* was divided by that of *GAPDH* to give the relative expression level of *Fbxo47* to *GAPDH.* Relative expression level of *Fbxo47* to *GAPDH* was normalized to 1 for a given P10 WT sample. Bar graph indicates mean with SD. Statistical significance was determined by t-test.

Figure 1F RNA was obtained from WT Embryonic ovaries (E12.5 to E18.5). qPCR was performed in triplicates or quadruplicates, and the average ddCt value was calculated for each cDNA sample. The expression level of *Fbxo47* was divided by that of *GAPDH* to give the relative expression level of *Fbxo47* to *GAPDH.* Relative expression level of *Fbxo47* to *GAPDH* was normalized to 1 for a given E12.5 WT sample. Bar graph indicates mean with SD.

Figure 3A Testis sections (P15) were obtained from non-tagged control (3 animals) and *Fbxo47-3FH* KI (3 animals). Number of seminiferous tubules that have HA+/ SYCP3+ cells was counted per the seminiferous tubules that have SYCP3+ spermatocyte cells (84, 85, 45 tubules for non-tagged control, 123, 90, 79 tubules for *Fbxo47-3FH* KI).

Figure 3D Testis sections (P18) were obtained from non-tagged control (3 animals) and *Fbxo47-3FH* KI (3 animals). Number of seminiferous tubules that have HA+/ H1t+ cells was counted per the seminiferous tubules that have H1t+ spermatocyte cells (52, 36, 18 tubules for non-tagged control; 15, 51, 36 tubules for *Fbxo47-3FH* KI).

Figure 4C Quantification of testes/body-weight ratio (mg/g) in *Fbxo47*+/- (8w; n=4) and *Fbxo47* KO (8w; n=10) mice. n: the number of animals examined for each genotype. Bar graph indicates mean with SD. Statistical significance was determined by t-test.

Figure 4I Cumulative number of pups born from *Fbxo47*+/- (n=4, all 6-week old at the start point of mating) and *Fbxo47* KO (n=4, all 6-week old at the start point of mating) females was counted for 18 weeks of breeding.

Figure 5A Quantification of the seminiferous tubules that have H1t+/SYCP3+ cells per the seminiferous tubules that have SYCP3+ spermatocyte cells in *Fbxo47* heterozygous (p18: 62, 61, 29 tubules/animal were counted from n= 3 animals; 8w: 135, 143, 45 tubules/animal were counted from n= 3 animals) and *Fbxo47* KO (p18: 105, 59, 141 tubules/animal were counted from n= 3 animals; 8w: 36, 55, 63, 64, 88, 69, 108 tubules/animal were counted from n= 7 animals) testes. n: the number of animals examined for each genotype. Bar graph indicates mean with SD. Statistical significance was determined by unpaired t-test. *p* = 0.0012 for *Fbxo47* heterozygous versus *Fbxo47* KO at P18.

Figure 5C Spermatocytes in the four developmental stages (leptotene, zygotene, pachytene, and diplotene(-like)) per total cells in meiotic prophase were quantified in WT (n=727 from one animal) and *Fbxo47* KO (n=659 from one animal) at P15, and in WT (n=561 from one animal) and *Fbxo47* KO (n=516 from one animal) at P18.

Figure 5F Spermatocytes in the four developmental stages (leptotene or zygotene, pachytene, and diplotene(-like)) per total cells in meiotic prophase were quantified in *Fbxo47*+/- (n=105 for OA -, 117 for OA+) and *Fbxo47* KO (n=140 for OA -, 117 for OA+).

Figure 5G Quantification of the seminiferous tubules that have TUNEL+ cells per total tubules in *Fbxo47*+/- (8w: n=3) and *Fbxo47* KO (8w: n=3) testes. Bar graph indicates mean with SD. Statistical significance was determined by t-test.

Figure 6C Numbers of RAD51 foci on SYCP3 axes were counted in WT and *Fbxo47* KO. Number of foci was indicated in the scatter plot with median. Statistical significance was determined by Mann-Whitney U-test.

Figure 6D Numbers of MSH4 foci on SYCP3 axes were counted in WT and *Fbxo47* KO. Number of foci was indicated in the scatter plot with median. Statistical significance was determined by Mann-Whitney U-test.

Figure 6E Numbers of MLH1 foci on SYCP3 axes were counted in *Fbxo47*+/- pachyene (n=15), or *Fbxo47* KO pachyene (n=13) and diplotene-like (n=13). Number of foci was indicated in the scatter plot with median. Statistical significance was determined by Mann-Whitney U-test.

Figure S3D

**Supplementary Figure 1.**
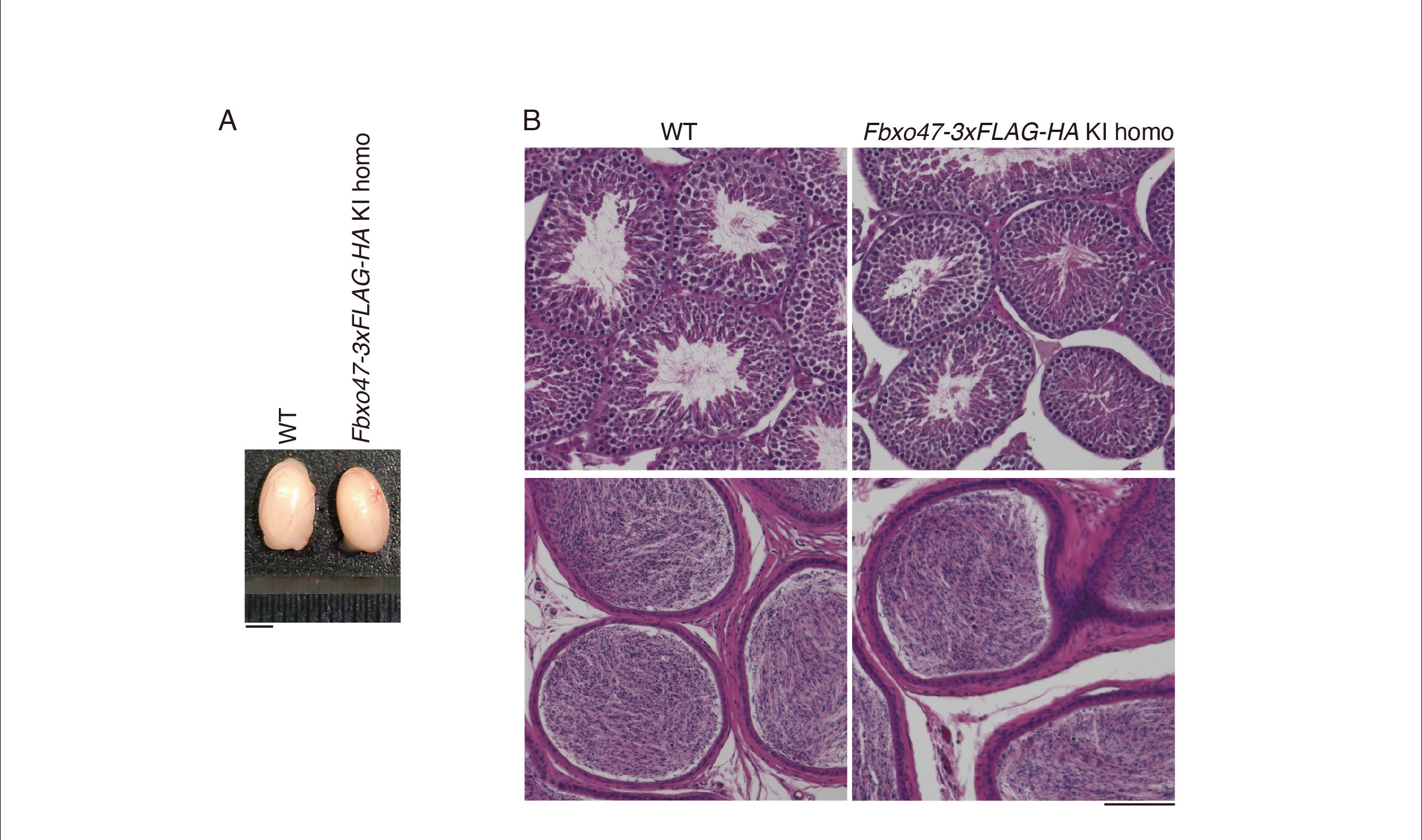
Generation of *Fbxo47-3xFLAG-HA* knock-in mice. **(A)** Testes from WT (no-tagged) and the *Fbxo47-3xFLAG-HA* KI homozygous mice (8-weeks old). Scale bar: 5 mm. **(B)** Hematoxylin and eosin staining of the testes (upper) and epididymis (lower) sections from WT (non-tagged control) and the *Fbxo47-3xFLAG-HA* KI homozygous testes (8-weeks old). Scale bar: 100 μm. Note that The FBOX47-3xFLAG-HA fusion protein was physiologically functional considering the normal fertility shown in homozygous male and female mice with the KI allele.

**Supplementary Figure 2.**
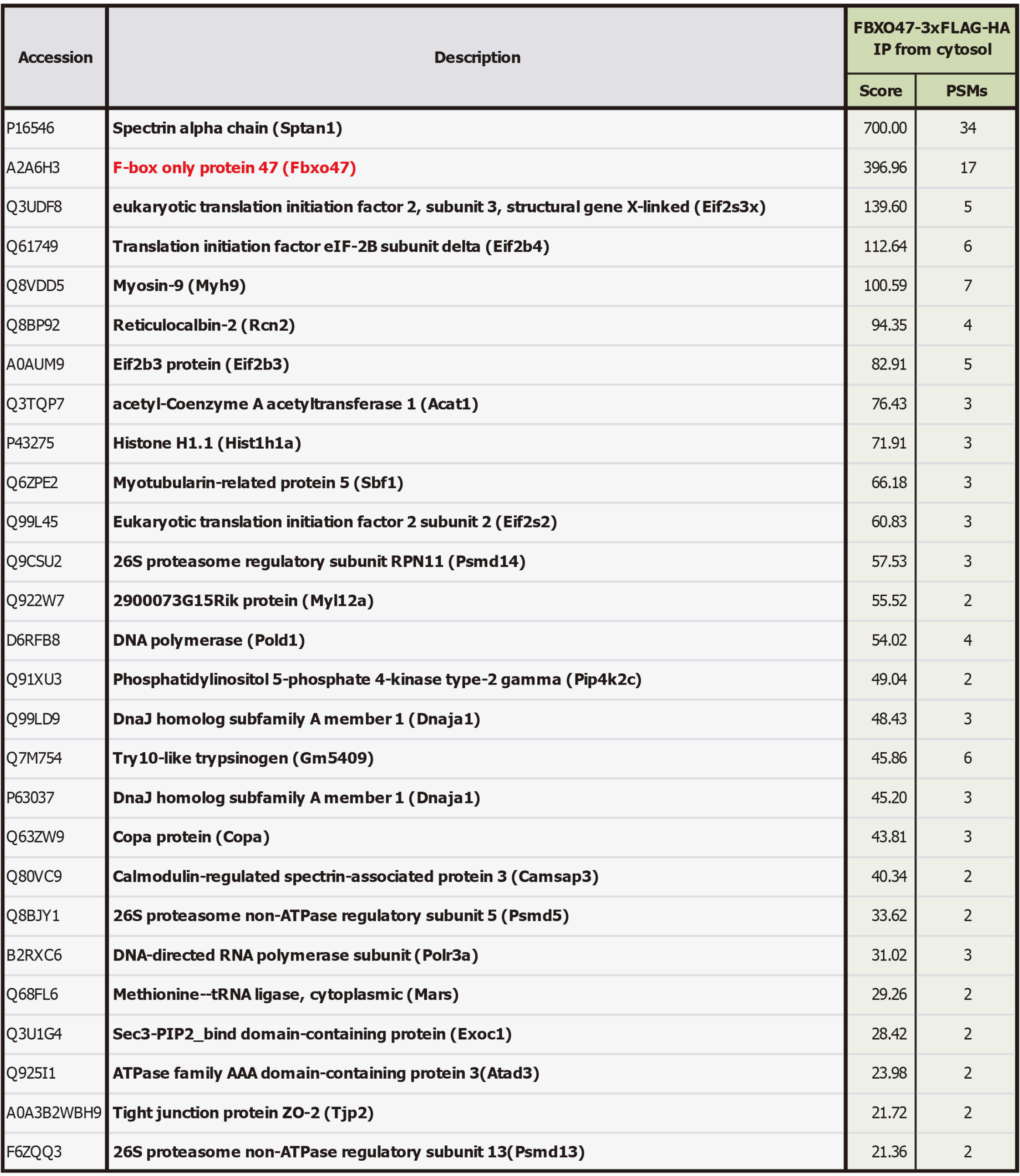
MS analyses of FBXO47 interacting factors in testis extracts. The immunoprecipitates (IP) from the cytosolic fraction of the testis extracts were subjected to liquid chromatography tandem-mass spectrometry (LC-MS/MS) analyses. The proteins identified by the LC-MS/MS analysis of FBXO47-IP are presented after excluding the proteins detected in the control IgG-IP. The proteins with more than 1 different peptide hits are listed with UniProt accession number, the number of peptide hits and Mascot scores.

**Supplementary Figure 3.**
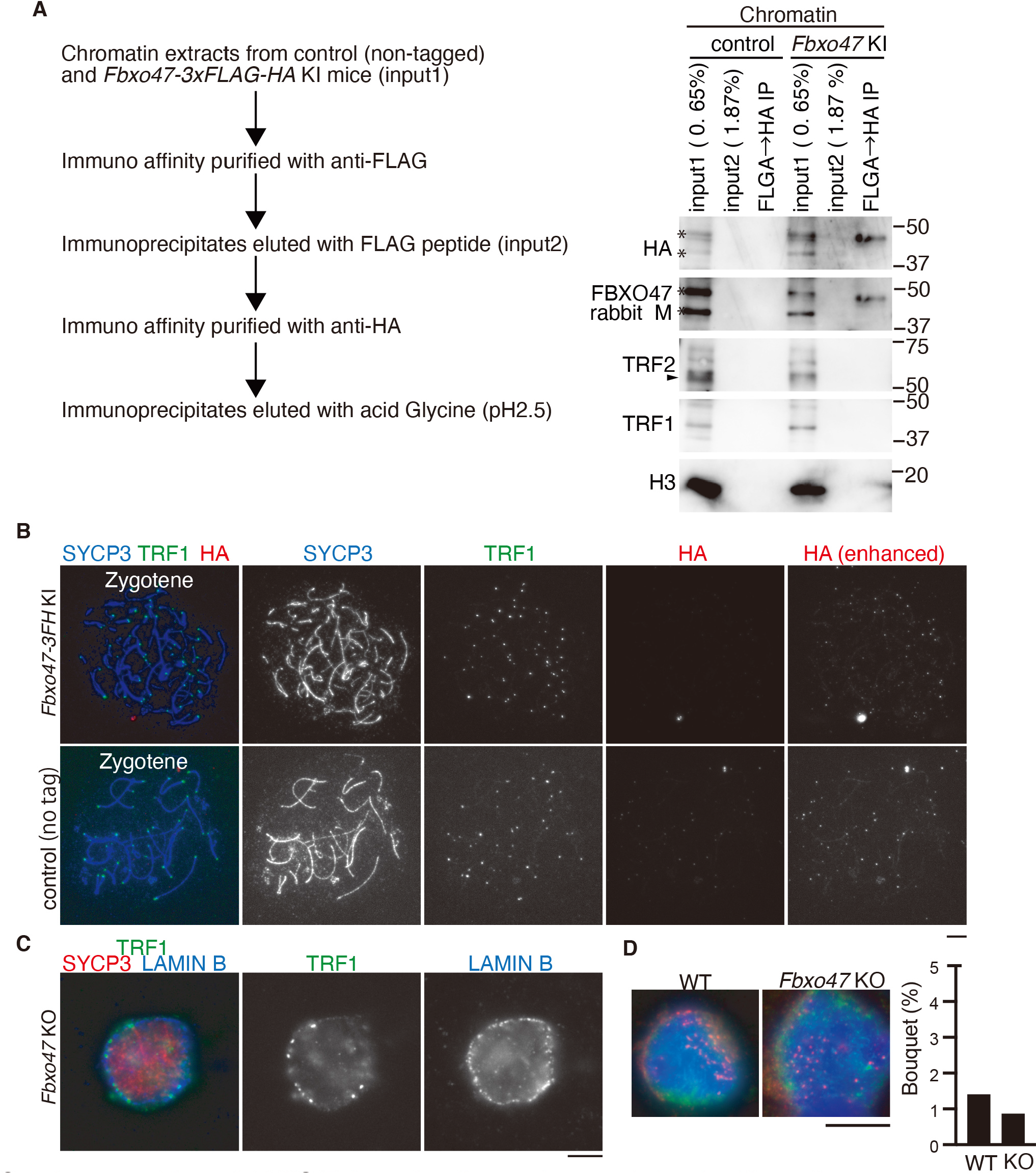
FBXO47 do not localize to telomeres. **(A)** Western blot showed immunoprecipitates from chromatin extracts of WT (non-tagged control) and *Fbxo47-3FH* KI mouse testes (from 139 and 148 animals at P14-19, respectively) after tandem affinity purifications using anti-FLAG and anti-HA antibodies. The same membrane was sequentially reblotted with different antibodies as indicated. *: non-specific band. Arrowhead: TRF2. Note that western blot did not detecte either TRF1 or TRF2 in the FBXO47 immunoprecipitate. **(B)** Chromosome spreads of *Fbxo47-3FH* KI and control (non-tagged) spermatocytes were immunostained as indicated. Images with enhanced contrast for HA color channel are shown. Scale bar: 5 μm. Note that FBXO47 did not show specific localization pattern to telomeres. We observed no more than background signals, even though contrast for HA images was enhanced. **(C)** Structurally-preserved nuclei of spermatocytes were prepared by squashing *Fbxo47* KO testis tubles, and immunostained for LAMIN-B, TRF1 and SYCP3. The image acquired at the equator of the spermatocyte nuclei is shown. Note that telomeres attachement to the nulear envelope was intact in *Fbxo47* KO spermatocytes. **(D)** The indicated spermatocyte nuclei were immunostained as indicated (Upper). Telomere clustering in wild-type (n=355) and *Fbxo47* KO (n=342) was scored at 12 day post-partum. The frequency of bouquet stage spermatocytes is shown (Bottom). Statistical significance is shown byN.S. *p* = 0.5025 (chi square-test).

**Supplementary Figure 4.**
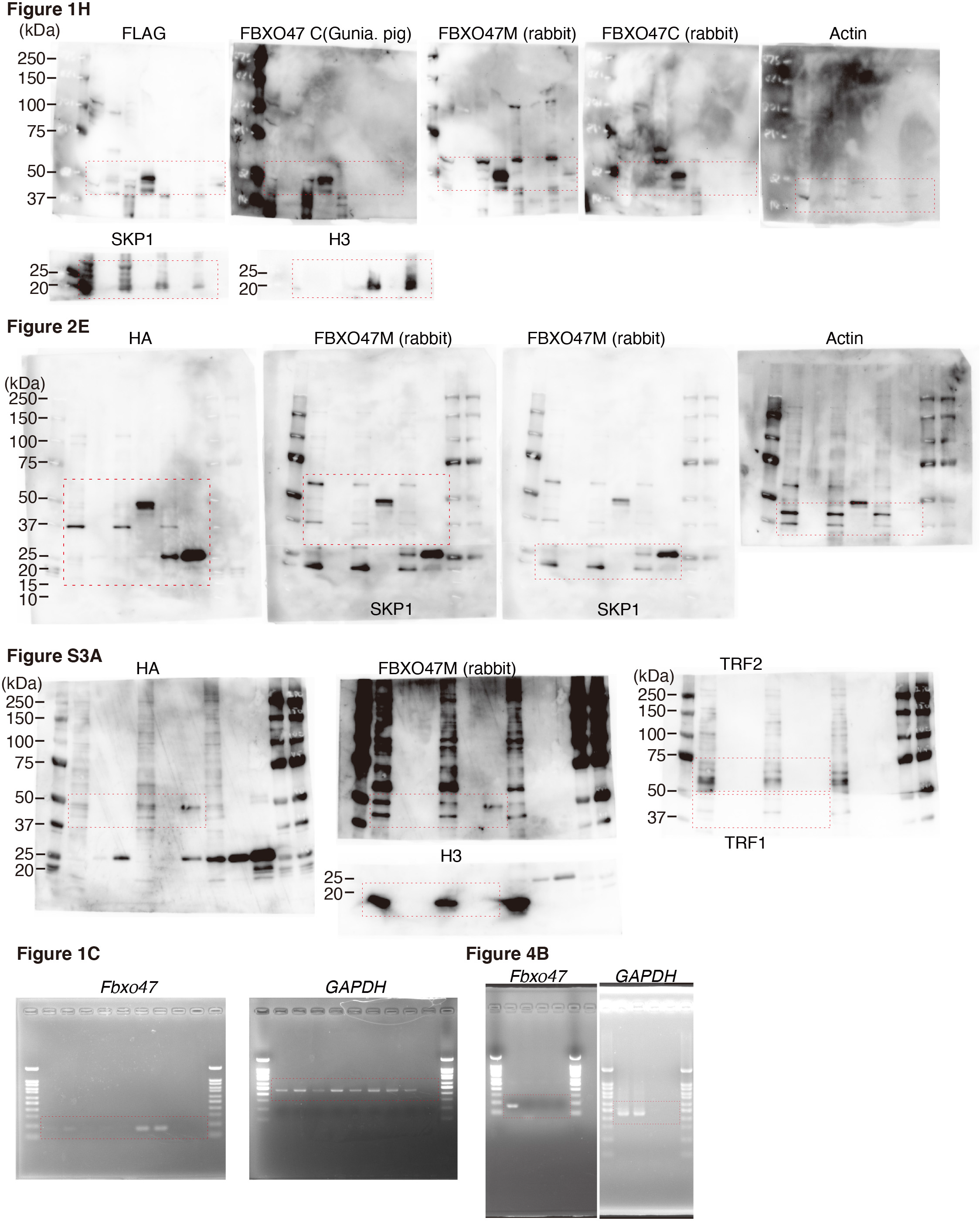
Uncropped images of gels and blots. Full-length / uncropped images of agarose gel (Fig1C, Fig4B) and immunoblots (Fig1H, Fig2E, Fig S3A) are shown. Immunoblotted membrane was sequentially reprobed with different antibodies. For SKP1, H3, TRF1 immunoblots, the same membrane was stripped, cut according to molecular weight marker and reprobed with different antibody, so that different proteins could be simultaneously probed with different antibodies. For Fig2E, the membrane was first immunoblotted with anti-HA antibody. After stripping, immunoblotted membrane was cut at a height of between 37 kDa and 25 kDa. Upper and lower membrane was immunoblotted with rabbit anti-FBXO47 middle region antibody and rabbit anti-SKP1 antibody, respectively. The membrane was combined and images were acquired sequentially at different exposure time.

## Notes

### Competing Interest Statement

The authors have declared no competing interest.

